# Acute, low-grade inflammation elevates white matter sensitivity to ischemia

**DOI:** 10.64898/2026.01.07.698147

**Authors:** Alexander G. Mellor, Glenn M. Harper, Eve E. Kelland, Robert Fern

## Abstract

Ischemic brain injuries, including stroke, are highly prevalent neurological diseases. Acute, low-grade inflammation, often resulting from peripheral infection, is a recognised risk factor. White matter axons, and their supporting glia, are susceptible to both ischemic and neuroinflammatory injury. The mechanisms behind ischemic and severe inflammatory injury in white matter are well established. However, the white matter response to acute, low-grade inflammation, and how this can interact with co-morbidities such as cerebral ischemia are yet to be examined. Here, we examine the response of white matter to acute, low-grade inflammation and the downstream effects this had on sensitivity of ischemia.

*Ex vivo* mouse optic nerve and corpus callosum, central white matter tracts, tolerated acute, low-grade inflammation (110-minutes, 0.1μg/ml LPS), with preserved action potential conduction, limited microglial activation and no axo-myelinic damage. Exposure to these conditions elevated indicators of stress in oligodendrocytes which, when exposed to ischemic conditions (30 minutes oxygen-glucose deprivation), impaired their ability to maintain healthy myelin. There was a resulting increase in axo-myelinic damage and functional decline associated with ischemic conditions compared to white matter that had not been pre-exposed to LPS. Microglial depletion or dampening microglial responses with the clinically available drug minocycline eliminated the elevated sensitivity of white matter to ischemic injury associated with acute, low-grade inflammation.

We demonstrate that acute, low-grade neuroinflammation primes white matter for heightened vulnerability to ischemic injury through microglial-mediated mechanisms. These findings provide a mechanistic explanation for the increased risk of ischemic brain injury observed during systemic inflammatory states and highlights microglial modulation as a clinical strategy to mitigate ischemic damage in susceptible patients.

## Introduction

Ischemic brain injuries, CNS damage caused by reduced cerebral blood flow^1^, are a prevalent neuropathology causing a high burden of death and disability globally^2–4^. Systemic infections and inflammatory conditions that trigger neuroinflammation increase the risk of ischemic brain injuries^5–14^. High infectious burden raises ischemic stroke risk by 1.4-fold^5^; hospitalization for infection elevates the likelihood of ischemic stroke and vascular dementia^6,7^; and periodontal disease increases stroke risk by 1.24-fold, particularly for lacunar strokes affecting white matter^12^. Despite these associations, the underlying mechanisms remain unclear.

White matter contains the myelinated axonal tracts in the CNS, where oligodendrocytes generate the myelin sheaths that ensheathe the large diameter axons responsible for rapid and efficient saltatory action potential transmission; provide metabolic support for these axons; and contribute to plasticity in the CNS^15^. These structures are highly vulnerable to ischemia and neuroinflammation, with white matter damage present in most ischemic brain injuries. For example, 80% of ischemic stroke patients present with white matter damage^16^. Myelin damage can lead to axonal degeneration and prolonged functional neurological deficits, and the extent of white matter injury strongly predicts disability^17,18^. Treatment for cerebral ischemia remains limited to risk factor control and restoring perfusion, with few CNS-specific or reparative therapies^16^. Significant levels of research has focused on neuroinflammation, particularly in inflammatory demyelinating diseases such as multiple sclerosis or progressive neurodegenerative diseases^19^. Modeling these diseases uses severe, long-term models, with examination weeks or months after initial insult with little investigation of early-stage damage, known to be an early pathology preceding demyelination^20^. The impact of acute, low-grade inflammation, which may cause subtle rather than overt injury, remains largely unexplored.

Ischemic and neuroinflammatory injuries share common mechanisms. White matter is highly susceptible to excitotoxicity via glutamatergic and purinergic receptors, leading to Ca^2+^-mediated oligodendrocyte and myelin damage^21–26^. Both conditions also induce oxidative stress^27–34^. Reactive astrocytes and microglia further contribute through excitotoxicity, reactive oxygen species (ROS) production and pro-inflammatory cytokine release during the acute phase^35–38^. These shared pathways make interactions between overlapping injury contexts, especially those with epidemiological links, an important area of study.

Modelling combined ischemia and low-grade neuroinflammation is complex due to spatial and temporal aspects of injury, variability and complexity of *in vivo* models. To address this, we used *ex vivo* white matter preparations from wild-type and transgenic mice, applying oxygen-glucose deprivation (OGD) to mimic ischemia and low-dose lipopolysaccharide (LPS) as a TLR-4-mediated inflammatory stimulus^39,40^. This approach enabled real-time assessment of white matter function via compound action potential (CAP) recordings and post-hoc microscopic analysis of cellular and subcellular injury.

Given the high number of ischemic brain injuries and the apparent risk of these caused by acute, low-grade neuroinflammation resulting from peripheral inflammation, there is potential for co-morbidities to interact and raise the risk or exacerbate injury, including in white matter. We therefore hypothesized that acute, low-grade neuroinflammation elevates the sensitivity of white matter to ischemia.

## Materials and methods

### Animals

Animal use complied with University of Plymouth ethical guidelines, the UK Animals (Scientific Procedures) Act 1986 and the ARRIVE guidelines. Mice were housed in individually ventilated cages under 12-hour light/dark cycles with food and water *ad libitum*. Adult (>4 months), mixed-gender CD-1 mice (Charles River, UK) and heterozygous PLP-GFP (B6;CBA-Tg(Plp1-EGFP)10Wmac/J, Jackson Laboratory, USA^41^) were used. PLP-GFP mice were genotyped using standard protocols.

### Reagents and drugs

Reagents were purchased from Fisher Scientific, UK unless specified. Solutions for *ex vivo* protocols were bubbled with 95% O_2_/5% CO_2_ or 95% N_2_/5% CO_2_ (both BOC, UK) and buffered to pH7.45. Lipopolysaccharide (*Escherichia coli*) was from Sigma-Aldrich, UK and Minocycline Hydrochloride from Bio-techne Tocris, UK. For microglial depletion, PLX5622 (Chemgood, USA) was incorporated into AIN-76A rodent diet (Research Diets Inc, USA) at 300 parts per million. Mice received PLX5622 diet, or matched PLX5622-free diet (AIN-76A), *ad libitum* for 21 days.

### *Ex vivo* preparations

#### CAP recordings

CAP recordings were performed as previously described^42^. Dissected mouse optic nerves (MONs) were maintained at 37±1°C in a perfusion chamber and superfused with oxygenated artificial CSF (aCSF; in mM: NaCl, 126; KCl, 3; NaH_2_PO_4_, 2; MgSO_4_, 2; CaCl_2_, 2; NaHCO_3_, 26; glucose, 10. pH 7.45). CAPs were evoked every 30 seconds using supramaximal square-wave, constant current pulses. (1-3mA, Iso Stim A320, World Precision Instruments). CAPs were differentially amplified and filtered (10,000X, 100-10,000Hz, CyberAmp 320, Axon Instruments or DP-311 Differential Amplifier, Warner Instruments) and digitised (Micro1401-3, Cambridge Electronic Design) and recorded using Signal Software version 7 (Cambridge Electronic Design). Area under curve (AUC) of the CAP trace was measured and values were normalised to a 100% baseline function (mean AUC of first 10 minutes) and 0% function (mean AUC of 5 minutes recording after MON crush).

#### Free floating *ex vivo* preparations

Rostral brain sections (250μm) were cut in ice-cold cutting solution (in mM: NaCl, 92; KCl, 2.5; NaH_2_PO_4_, 1.2; MgSO_4_, 2; CaCl_2_, 2; NaHCO_3_, 30; Glucose, 25; HEPES, 20; Na Pyruvate, 3; Thiourea, 2; Na Ascorbate, 5. pH 7.4) to expose the mouse corpus callosum (MCC), hemi-sectioned and placed in aCSF at 37±1°C as matched pairs. MONs from the same brains were also collected and maintained similarly.

#### Oxygen-glucose deprivation and lipopolysaccharide exposure

*Ex* vivo experimental time courses are shown in supplementary figure 1. After recovery and baseline recordings in either aCSF or 0.1μg/ml LPS (in aCSF), preparations were exposed to: (1) 110-minute control (in aCSF), (2) 30 minutes OGD with nitrogenated, glucose-free aCSF (in mM: NaCl, 126; KCl, 3; NaH_2_PO_4_, 2; MgSO_4_, 2; CaCl_2_, 2; NaHCO_3_, 26; sucrose, 10. pH 7.45) followed by a 70-minute recovery, (3) 0.1μg/ml LPS (in aCSF) or (4) combined 30 minutes OGD and 0.1μg/ml LPS. Preparations were fixed in 4% paraformaldehyde (PBS, pH 7.4) for fluorescent imaging or immunohistochemistry or 2.5% glutaraldehyde (0.1M sodium cacodylate, pH 7.2) for electron microscopy.

#### Fluorescent imaging

Images were acquired using a confocal microscope (Leica SPE and LasX software, Leica, Germany) and processed using ImageJ (NIH, USA).

#### Vital myelin imaging

Fixed *ex vivo* preparations were stained using FluoroMyelin™ Red (FMR) Fluorescent Myelin stain (ThermoFisher, 1:300 in PBS) for 60 minutes at room temperature, washed and mounted in Flouromount Aqueous mounting medium (Sigma-Aldrich).

#### PLP-GFP imaging

PLP-GFP tissue was stained with DAPI (FluoroPure Grade, Invitrogen, 1:10,000 in PBS, pH7.4), washed (3x5 mins, PBS), mounted and imaged.

#### Transmission Electron Microscopy

Transmission Electron Microscopy (TEM) was performed as previously described^42^. MONs were fixed in 2.5% glutaraldehyde (0.1M sodium cacodylate, pH7.2), post-fixed in 1% osmium tetroxide, dehydrated in graded ethanol (30-100%) and infiltrated with increasing concentrations of epoxy resin, then polymerised at 60°C overnight. Ultra-thin sections (80nm) were mounted on copper grids, counterstained with uranyl acetate or neodymium acetate and lead citrate and imaged using a JEOL JEM 1400 TEM with Gatan Orius SC200W or Rio cameras (DigitalMicrograph software).

### Immunohistochemistry

#### Antibodies

Primary antibodies were GFAP (Astrocytes, Rat IgG2a monoclonal anti-GFAP, abcam, ab279291, 1:500), Iba1 (Microglia, Rat IgG2a monoclonal anti-Iba1, abcam, ab283346, 1:1000), pan-NF-H (axons, Chicken IgY polyclonal anti-Neurofilament Heavy Polypeptide, abcam, ab4680, 1:1000) and SMI-32 (non-phosphorylated-NF-H, Mouse IgG1 monoclonal anti-neurofilament H non-phosphorylated antibody, Calbiochem, NE1023, 1:1000).

Secondary antibodies were AlexaFluor® 488 Goat Anti-Rat IgG H&L (Abcam, ab150157, 1:1000), AlexaFluor® 647 Goat Anti-Mouse IgG H&L (Abcam, ab150115, 1:1000) and Alexa Fluor® 568 Donkey anti-Chicken IgY H&L (Invitrogen, A78950, 1:1000).

No cross-reactivity was observed in secondary-only controls.

#### Protocol

Fixed MONs were cryoprotected (30% sucrose in PBS, 48 hours), embedded (OCT compound, CellPath) and cryosectioned (10µm). Cryosections were washed (3x5mins, TBS), quenched (3% H_2_O_2_, 10% Methanol in TBS) washed again (3x5 mins, TBS), blocked (4% Normal Goat Serum (abcam), 2% Normal Horse Serum (abcam), 0.1% Triton X-100 in TBS) at room temperature and incubated with primary antibodies (in 1% NGS, 0.1% Triton X-100 in TBS) at 4°C for 24 hours. After washing (15 mins, 0.1% Triton X-100 in TBS, 15 mins, TBS), secondary antibodies were applied (in 1% NGS, 1% NHS, 0.1% Triton X-100 in TBS) at 4°C for 24 hours. Cryosections were washed (15 mins, 0.1% Triton X-100 in TBS, 15 mins, TBS) and mounted (FluoroMount-G with DAPI, Invitrogen).

### Image analysis

All analyses were performed using ImageJ (NIH, USA). Regions of interest were selected freehand, and area/distance measurements were taken using built-in tools. Microglial morphology and EM images were analysed blind.

#### Vital myelin imaging and fluorescent image analysis

Fluorescence intensity was measured as the mean grey value across the MCC/MON. Cell and axon densities were quantified as GFP/GFAP/Iba1 or NF-H/SMI-32-positive counts per area. Astrocyte morphology was assessed via binary pixel analysis (GFAP-positive/negative). Microglial morphology was classified as: ramified: fine, branched processes; bushy: truncated processes, swollen soma; ameboid: rounded soma, no processes, motile: two thin bipolar processes^42^.

#### Electron microscopy analysis

Electron microscopy analysis is detailed in Supplementary Figure 2. Axon and fibre diameters were measured by fitting axons as idealised circles. G-ratio is the ratio between axon diameter and total fibre diameter. In still-compact myelin, myelin thickness was measured as the distance between the inner tongue to outer lamella and periodicity the myelin thickness divided by the lamellae count. Semi-quantitate myelin pathology scoring of focal myelin pathologies was defined as: (1) no pathology, (2) slight splitting between layers, (3) several points of decompaction and loss of uniformity, and (4) severe decompaction and loss of integrity.

### Statistical analysis

Statistical testing was performed using GraphPad PRISM v9.5.1. Statistical significance was set at *P*<0.05. Data is shown as individual datapoints ± SEM (unpaired) or paired datapoints. Sample sizes were based on power calculations using prior variability^42^. Outliers were assessed using Grubb’s test; none were excluded. Normality was tested with Shapiro-Wilk’s test (if sample size allowed); otherwise, normality was assumed. Normal data was analysed using paired/unpaired one-way t-tests or one-way ANOVAs with Holm-Šídák’s or Bonferroni’s post hoc tests and non-normal data using a Kruskal-Wallis test with Dunn’s post hoc test. The timecourse of the AUC was analysed using a two-way repeated measures ANOVA with Holm-Šídák’s post-hoc test. Significance denotation: ns=non-significant (P>0.05), *=P≤0.05, **=P≤0.01, ***=P≤0.001, ****=P≤0.0001.

### Data availability

The data that supports the findings of this study are available from the corresponding author upon reasonable request.

## Results

### Acute, low-dose LPS pre-application causes a reduction in the functional recovery of white matter following OGD

We first employed CAP recordings in the MON to assess white matter function (Fig 1), with acute, low-grade inflammation induced with 0.1μg/ml LPS throughout the recovery and recording period and ischemia induced with 30 minutes OGD + 70 minutes recovery.

**Figure 1:**
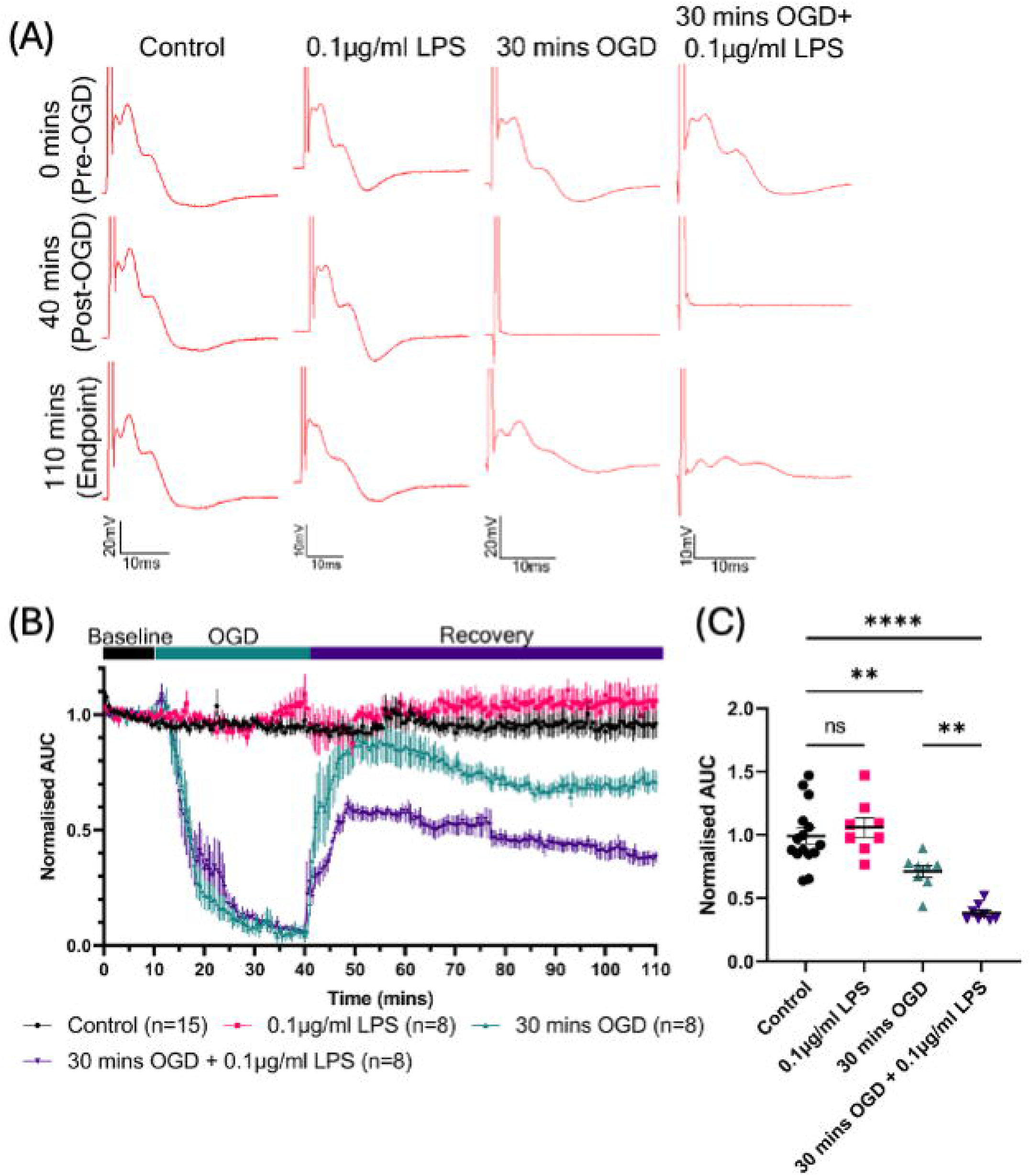
0.1µg/ml LPS pre-treatment caused no loss of optic nerve function but elevated functional loss following 30 minutes of OGD. (A) Representative CAP traces at 0 mins (baseline, pre-30 mins OGD), 40 mins (post-30 mins OGD), and 110 mins (endpoint, post-70 mins recovery). (B) Normalised area under the curve of CAP recordings over time in control optic nerves and optic nerves treated with 0.1μg/ml LPS, 30 mins OGD or both (Two-way repeated measures ANOVA with Holm-Šídák’s post hoc multiple comparisons). (C) Normalised area under curve of CAP recordings at 110 mins (endpoint) following 0.1μg/ml LPS, 30 mins OGD or both (One-way ANOVA with Holm-Šídák’s post hoc multiple comparisons).

The AUC over time showed significant differences between groups (Fig. 1B, *P*<0.0001). No significant differences were observed between control and 0.1μg/ml LPS pre-exposed MONs (*P>*0.05, all time points). Post-reperfusion, MONs exposed to 30 minutes OGD showed reduced recovery compared to controls (*P<*0.05, 68.5-110 mins), with a further reduction following 30 minutes OGD + 0.1μg/ml LPS pre-application (*P*<0.05, 51-110 mins).

At endpoint, AUCs differed significantly (Fig. 1C, *P*<0.0001). 0.1µg/ml LPS pre-application alone had no significant effect on endpoint AUC compared to control (Control=0.99±0.06, 0.1µg/ml LPS=1.06±0.08, *P*=0.4261). 30 minutes OGD significantly reduced the endpoint AUC (30 minutes OGD=0.71±0.05, *P=*0.0058) and 30 minutes OGD + 0.1µg/ml LPS pre-application further reduced the AUC at endpoint (30 mins OGD + 0.1μg/ml LPS=0.38±0.03, *P*<0.0001 vs. control, *P*=0.0058 vs. 30 mins OGD).

These AUC changes suggest that while low-grade inflammation alone does not impair white matter function acutely, it worsens recovery following ischemia.

### Combined low-dose LPS pre-application and OGD increases acute injury to myelin and their underlying axons

To assess structural damage underlying functional changes in the MON, we examined acute myelin and axonal injury.

Vital myelin imaging using FMR showed no significant changes in fluorescence in the MCC or MON following 0.1µg/ml LPS alone (MCC: Control=100.2±5.79, 0.1μg/ml LPS=104.5±9.53, *P*=0.2921. MON: Control=126.4±7.83, 0.1μg/ml LPS=130.7±6.71, *P*=0.0611) (Fig. 2A, F). 30 minutes OGD significantly reduced fluorescence in both the FMR-stained MCC and MON (MCC: Control=111.7±5.73, 30 minutes OGD=97.69±5.06, *P*=0.0021. MON: Control=135.3±13.56, 30 minutes OGD=101.0±6.11, *P*=0.0079) (Fig. 2B, F). 30 minutes OGD + 0.1µg/ml LPS pre-application significantly reduced fluorescence in the FMR-stained MCC and MON (MCC: Control=109.9±7.33, 30 minutes OGD + 0.1μg/ml LPS=75.75±9.00, *P*<0.0001. MON: Control=124.2±7.64, 30 minutes OGD + 0.1μg/ml LPS=80.21±9.77, *P*=0.0003) (Fig. 2C, F). Compared to 30 minutes OGD alone, 30 minutes OGD + 0.1µg/ml LPS pre-application significantly reduced fluorescence in FMR-stained MCCs and MONs (MCC: 30 minutes OGD=133.2±8.51, 30 minutes OGD + 0.1μg/ml LPS=115.3±7.97, *P*=0.0137. MON: 30 minutes OGD=172.5±14.09, 30 minutes OGD + 0.1μg/ml LPS=138.4±17.74, *P*=0.0187) (Fig. 2D, F).

**Figure 2:**
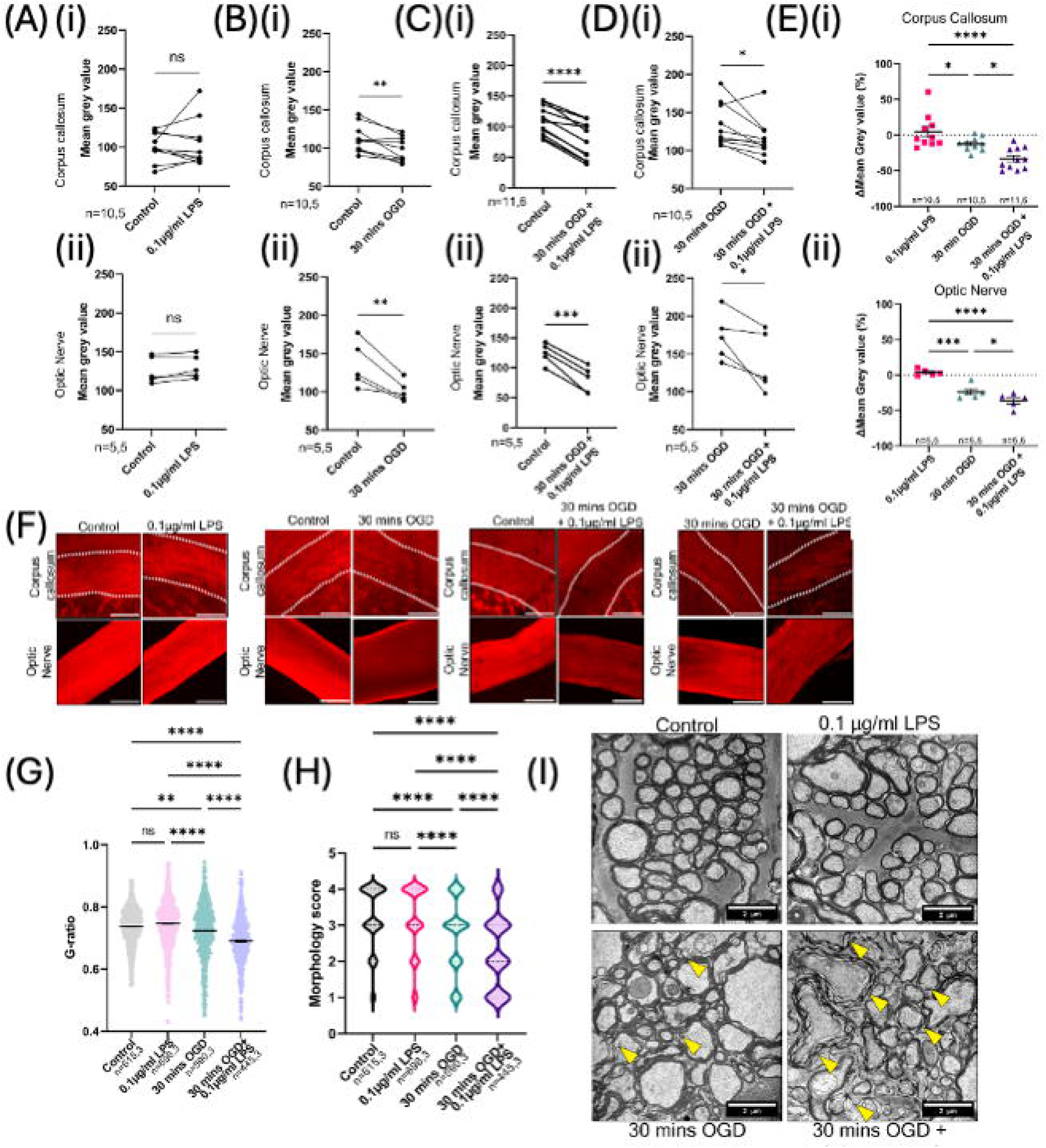
0.1μg/ml LPS pre-treatment does not cause acute myelin injury but elevates myelin injury combined with 30 minutes OGD. (A-D) Mean grey value of paired, FMR-stained corpus callosum hemi-sections **(i)** and optic nerves **(ii)** (paired t-test). **(E)** The change in mean grey value of FMR-stained corpus callosum hemi-sections **(i)** and optic nerves **(ii)** compared to control (One-way ANOVA with Holm-Šídák’s post hoc multiple comparisons). **(F)** 10X paired FMR-stained corpus callosum (dotted line) and optic nerves. **(G)** G-ratios of optic nerve axons following 0.1μg/ml LPS, 30 mins OGD or both (One-way ANOVA with Bonferroni’s post-hoc multiple comparison). **(H)** Myelin pathology scores of optic nerve myelin sheaths following 0.1μg/ml LPS, 30 mins OGD or both (Kruskal-Wallis test with Dunn’s post hoc multiple comparison). **(I)** 10,000X representative micrographs of the optic nerve; focal decompaction sites indicated by yellow arrows.

Percentage change relative to controls were significantly different in the MCC (*P*<0.0001) and MON (*P*<0.0001) (Fig. 2E). In both the MCC and MON, combined 30 minutes OGD + 0.1μg/ml LPS pre-application caused a significantly greater percentage increase compared to 30 minutes OGD alone (MCC: 30 minutes OGD=12.17±2.99%, 30 minutes OGD + 0.1μg/ml LPS=33.22±4.55%, *P*=0.0169. MON: 30 minutes OGD=23.9±4.52%, 30 minutes OGD + 0.1μg/ml LPS=36.19±4.65%, P=0.0459).

We validated the observations of acute myelin injury using TEM. G-ratio measurements in the MON showed significant differences (*P*<0.0001) (Fig. 2G). 0.1μg/ml LPS alone had no significant effects on the G-ratio (Control=0.74±0.002, 0.1μg/ml LPS=0.75±0.002, *P*=0.1231) but 30 minutes OGD significantly reduced the G-ratio (30 minutes OGD=0.72±0.003, *P*=0.0075). A further significant G-ratio decrease was seen following 30 minutes OGD + 0.1μg/ml LPS pre-application compared to control and 30 minutes OGD (30 minutes OGD + 0.1μg/ml LPS=0.69±0.004, vs. control *P*<0.0001, vs 30 minutes OGD *P*<0.0001).

Semi-quantitative analysis of the appearance of focal myelin decompactions and myelin integrity via pathology scoring confirmed significant differences in myelin pathology (*P*<0.0001) (Fig. 2H, I). No difference in pathology score was seen in control and 0.1μg/ml LPS pre-exposed MONs (Control=1.78±0.04, 0.1μg/ml LPS=1.83±0.04, *P*>0.9999) but 30 minutes OGD induced decompactions and losses of myelin integrity, significantly increasing the pathology score (30 minutes OGD=2.29±0.04, *P*<0.0001). Combined 30 minutes OGD + 0.1µg/ml LPS pre-application caused a further increase in pathology score compared to control and 30 minutes OGD MONs (30 minutes OGD + 0.1μg/ml LPS=2.78±0.05, vs. control *P*<0.0001, vs. 30 mins OGD *P*<0.0001).

We next assessed the axonal compartment of the MON. TEM identified differences in the number of axons in the MON (*P*=0.0393) (Fig. 3Ai), but post hoc testing did not identify which changes were significant (Control=0.45±0.05 axons/µm^2^, 0.1μg/ml LPS=0.46±0.03 axons/µm^2^, 30 minutes OGD=0.36±0.03 axons/µm^2^, 30 minutes OGD=0.28±0.04 axons/µm^2^, all comparisons, *P*>0.05). Staining for axons with heavy-chain neurofilament (Fig. 3Aii, E) showed no changes axonal density (controls=115596±13669 axons/mm^2^, 0.1µg/ml LPS=104597±4059 axons/mm^2^, 30 minutes OGD=113190±7235 axons/mm^2^, 30 minutes OGD + 0.1µg/ml LPS=99514±13699 axons/mm^2^, *P*=0.6893). We next assessed levels of SMI-32, an antibody against dephosphorylated neurofilaments, which can be used to detect axonal damage following acute damage to white matter axons (Sup. Fig. 3).

**Figure 3:**
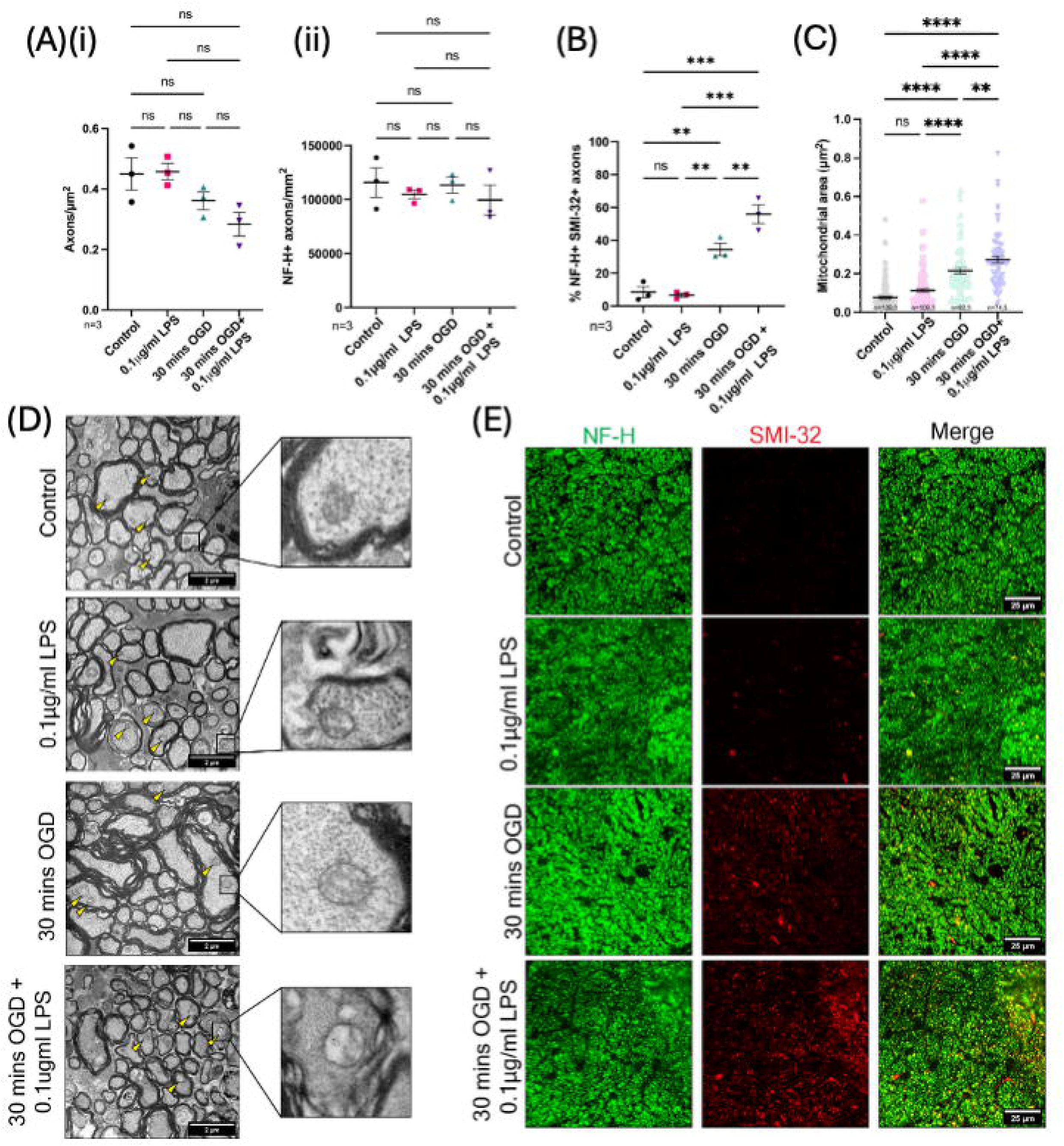
0.1μg/ml LPS does not cause axonal injury but elevates myelin injury when combined with 30 minutes OGD. **(A)** Axonal density in optic nerves following 0.1μg/ml LPS, 30 mins OGD or both calculated by both TEM **(i)** and NF-H immunohistochemistry **(ii)** (One-way ANOVA). **(B)** % of NF-H+ optic nerve axons positive for SMI-32 following 0.1μg/ml LPS, 30 mins OGD or both (One-way ANOVA with Holm-Šídák’s post hoc multiple comparisons). **(C)** Mitochondrial area of optic nerve axons following 0.1μg/ml LPS, 30 mins OGD or both (One-way ANOVA with Bonferroni’s post-hoc multiple comparison). **(D)** 10,000X representative micrographs of the optic nerve showing axonal mitochondria (yellow arrows, expansions). **(E)** 63X representative images of optic nerve axons immunostained for NH-H and SMI-32.

The percentage of axons positive for SMI-32 showed significant differences (*P*<0.0001) (Fig. 3B, E). The percentage of axons positive for SMI-32 did not change following 0.1μg/ml LPS (Control=8.52±3.29%, 0.1μg/ml LPS=6.72±1.22%, *P*=0.7481). 30 minutes OGD increased the percentage of axons positive for SMI-32 compared to control MONs (30 minutes OGD=34.44±3.77%, *P*=0.0041), further increasing following 30 minutes OGD + 0.1μg/ml LPS pre-exposure (30 minutes OGD + 0.1μg/ml LPS=55.86±5.67%, *P*=0.0084).

As mitochondrial swelling is a feature of ischemic injury in white matter axons^42^, we examined the area of axonal mitochondria in MON axons (Fig. 3C, D), showing a difference in mitochondrial areas between groups (*P*<0.0001). 0.1µg/ml LPS did not induce a significant change in axonal mitochondrial area (Control=0.08±0.01um^2^, 0.1μg/ml LPS=0.11±0.01µm^2^, *P*=0.0688). Following 30 minutes OGD, the area of axonal mitochondria was significantly increased (30 minutes OGD=0.22±0.02µm^2^, *P*<0.0001) and further increased following 30 minutes OGD + 0.1μg/ml LPS (30 minutes OGD + 0.1μg/ml LPS=0.27±0.02µm^2^, vs controls *P*<0.0001, vs 30 minutes OGD *P*=0.0084).

Low-grade inflammation alone does not acutely damage the axo-myelinic unit but exacerbates injury following ischemia, consistent with functional deficits.

### Oligodendrocytes are stressed under acute, low-dose LPS application

Given the identification of an elevated axo-myelinic injury following combined ischemia and low-grade inflammation, we next examined damage to oligodendrocytes, the myelinating cells of the CNS, in PLP-GFP mice in which all oligodendrocytes express green-fluorescent protein (GFP).

We observed no loss of oligodendrocyte numbers in the MCC or MON from PLP-GFP mice (Fig. 4A, Sup. Fig. 4.). Percentage changes in the cell density showed no significant differences in the MCC and MON (MCC: 0.1µg/ml LPS=1.66±5.89%, 30 minutes OGD=-8.06±10.97%, 30 minutes OGD + 0.1µg/ml LPS=4.08±6.33%, *P*=0.7322. ON: 0.1μg/ml LPS=-12.93±24.22%, 30 minutes OGD=-11.93±15.67%, 30 minutes OGD + 0.1µg/ml LPS=-11.57±16.54%, *P*=0.9987).

**Figure 4:**
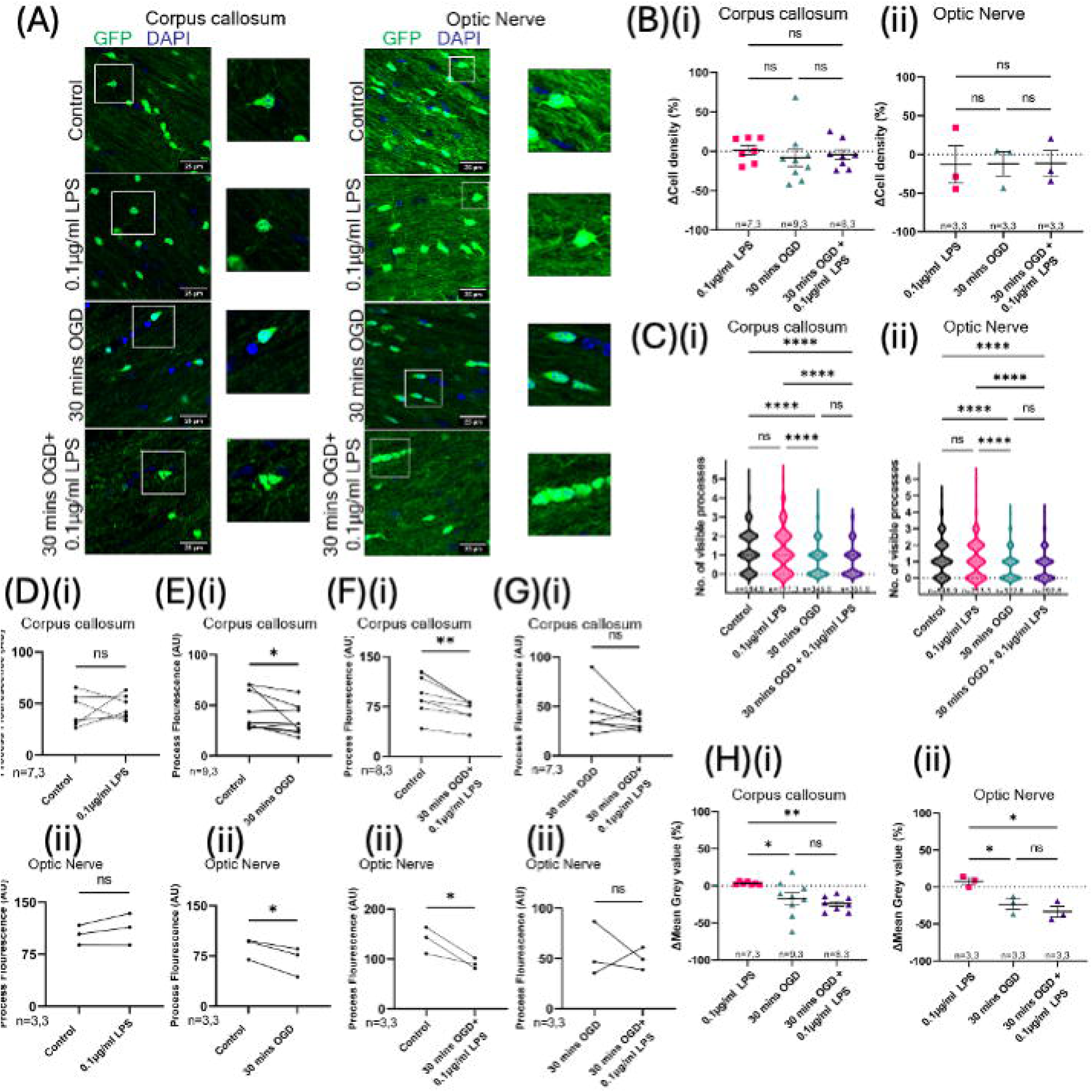
0.1μg/ml LPS causes no oligodendrocyte death or process damage and does not elevate process damage induced by 30 minutes OGD. **(A)** 63X images of GFP+ oligodendrocytes in the corpus callosum **(i)** and optic nerve **(ii)** following 0.1μg/ml LPS, 30 mins OGD or combined treatment. **(B)** The change in oligodendrocyte density in the corpus callosum **(i)** and optic nerve **(ii)**, relative to controls, following 0.1μg/ml LPS, 30 mins OGD or combined (One-way ANOVA with Holm-Šídák’s post hoc multiple comparisons). **(C)** Number of visible oligodendrocyte processes in the corpus callosum **(i)** and optic nerve **(ii)** following 0.1μg/ml LPS, 30 mins OGD or combined (One-way ANOVA with Bonferroni’s post-hoc multiple comparison). **(D-G)** Process fluorescence in paired corpus callosum **(i)** and optic nerves **(ii)** following treatment with 0.1μg/ml LPS, 30 mins OGD or combined (paired t-test). **(H)** The change in GFP mean grey value in the corpus callosum **(i)** and optic nerve **(ii)**, relative to controls, following 0.1μg/ml LPS, 30 mins OGD or combined (One-way ANOVA with Holm-Šídák’s post hoc multiple comparisons).

Whilst oligodendrocyte numbers remined unchanged, we looked at visible process loss from oligodendrocytes, which showed significant differences in both the MCC and MON (MCC *P*<0.0001, MON, *P*<0.0001) (Fig. 4A, C). 0.1μg/ml LPS caused no visible processes loss (MCC: Control=1.25±0.04, 0.1μg/ml LPS=1.25±0.09, *P*>0.9999. MON: Control=1.27±0.05, 0.1μg/ml LPS=1.18±0.08, *P*>0.9999). 30 minutes caused a significant loss of processes in MCCs and MONs (MCC: 30 minutes OGD=0.76±0.04, *P*<0.000. MON: 30 minutes OGD=0.74±0.05, *P*<0.0001). 30 minutes OGD + 0.1μg/ml LPS pre-application caused oligodendrocytes to lose processes in both the MCC and MON (MCC: 30 minutes OGD + 0.1μg/ml LPS= to 0.73±0.04, *P*<0.0001. ON: 30 mins OGD + 0.1μg/ml LPS=0.77±0.05, *P*<0.0001), but this loss was not elevated compared to 30 minutes OGD alone (MCC: *P*>0.9999. MON: *P*>0.9999).

Visible processes in PLP-GFP mice do not fully encapsulate the complexity of oligodendrocyte processes, hence we used a loss of non-somatic GFP as a surrogate for the loss of less visible processes. 0.1μg/ml LPS caused no loss of process fluorescence in the MCC or MON (MCC: Control=42.76±5.7, 0.1μg/ml LPS=46.21±4.27, *P*=0.3044. MON: Control=103.2±8.24, 0.1μg/ml LPS=111.9±13.22, *P*=0.1102) (Fig 4D). 30 minutes OGD caused a loss of process intensity in the MCC and MON (Fig 4E) (MCC: Control=44.18±6.33, 30 minutes OGD=34.87±4.82, *P*=0.0444. MON: Control=87.97±9.34, 30 minutes OGD=68.69±12.61, *P*=0.0174). Process intensity was decreased following 30 minutes OGD + 0.1μg/ml LPS pre-application (Fig. 4F) (MCC: Control=93.93±10.48, 30 minutes OGD + 0.1μg/ml LPS=69.66±6.09, *P*=0.0019. MON: Control=139.1±15.29, 0.1μg/ml LPS=90.91±6.02, *P*=0.0355), but this process intensity loss was not greater than following 30 minutes OGD (Fig. 4G) (MCC: 30 minutes OGD=44.87±8.53, 30 minutes OGD + 0.1μg/ml LPS=34.74±2.76, *P*=0.1085. MON: 30 minutes OGD=56.33±15.48, 30 minutes OGD + 0.1μg/ml LPS=49.65±6.41, *P*=0.3746). The percentage change in process intensity relative to controls differed between groups in both the MCC and MON (MCC *P*=0.009, MON *P*=0.0091) (Fig. 4H), but the percentage loss was not different following 30 minutes OGD or 30 minutes OGD + 0.1μg/ml LPS pre-application (MCC: 30 minutes OGD=16.9±7.96%, 30 minutes OGD + 0.1μg/ml LPS=24.08±3.52%, *P*=0.3620. MON: 30 minutes OGD=23.34±6.97%, 30 minutes OGD + 0.1μg/ml LPS=33.33±7.36%, *P*=0.3091).

As a measure of mitochondrial and oxidative stress, we measured whether mitochondria within MON oligodendrocytes swell following 0.1µg/ml LPS pre-application, 30 minutes OGD or both, showing significant changes in mitochondrial area following 0.1µg/ml LPS, 30 minutes OGD or combined 30 minutes OGD and 0.1µg/ml LPS pre-application (*P*<0.0001) (Fig. 5A, B). Mitochondria swelled following 0.1μg/ml LPS (Control=0.042±0.002µm^2^, 0.1μg/ml LPS=0.063±0.003µm^2^, *P*<0.0001), to a similar level to swelling seen following 30 minutes OGD (30 minutes OGD=0.06±0.004µm^2^, vs Control *P*<0.0001, vs 0.1μg/ml LPS P>0.9999). 30 minutes + 0.1µg/ml LPS pre-application caused oligodendrocyte mitochondria to swell further compared to both controls and 30 minutes OGD alone (30 minutes OGD + 0.1μg/ml LPS=0.08±0.004µm^2^, vs control *P*<0.0001, vs 30 minutes OGD *P*<0.0001).

**Figure 5:**
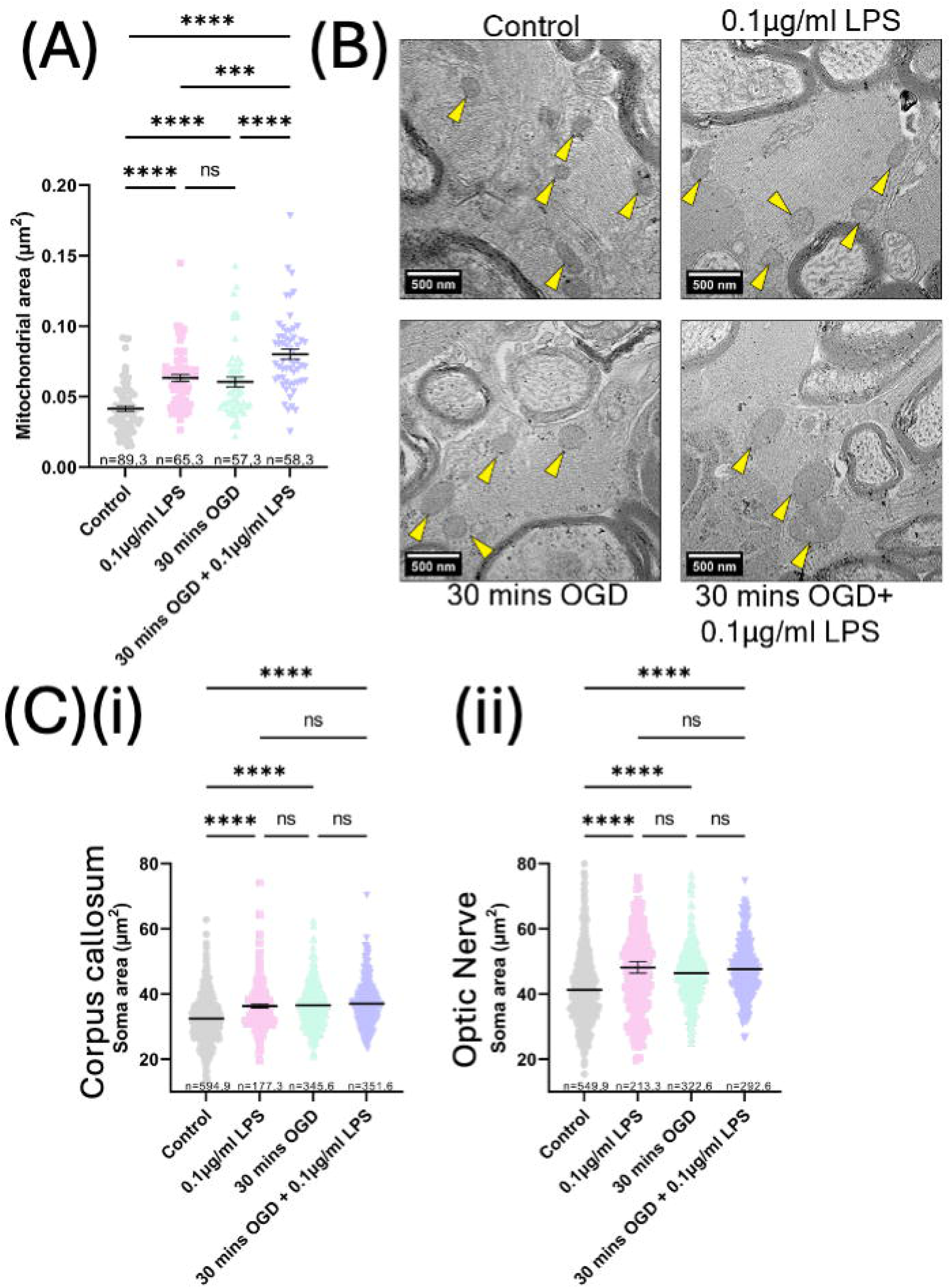
0.1μg/ml LPS causes oligodendrocyte mitochondria and soma swelling to similar levels as 30 minutes OGD. **(A)** The mitochondrial area of optic nerve oligodendrocyte mitochondria following 0.1μg/ml LPS, 30 mins OGD or combined treatment (One-way ANOVA with Bonferroni’s post-hoc multiple comparison). **(B)** 10,000X micrographs of optic nerve oligodendrocyte mitochondria (yellow arrows) following 0.1μg/ml LPS, 30 mins OGD or both. **(C)** Oligodendrocyte soma areas in the corpus callosum **(i)** and optic nerves **(ii)** following 0.1μg/ml LPS, 30 mins OGD or both (One-way ANOVA with Bonferroni’s post-hoc multiple comparison).

Oligodendrocyte swelling is a feature of ischemic stroke^43^, hence we looked at whether the soma of oligodendrocytes had exacerbated swelling following 0.1μg/ml LPS pre-application and 30 minutes OGD. Oligodendrocytes in the MCC (*P*<0.0001) and MON (*P*<0.0001) both underwent significant changes in the soma area following 0.1µg/ml LPS, 30 minutes OGD or both (Fig. 5C). 0.1μg/ml LPS alone caused oligodendrocyte soma swelling (MCC: Control=32.44±0.55µm^2^, 0.1μg/ml LPS=36.27±0.57µm^2^, *P*<0.0001. MON: Control=41.23±0.46µm^2^, 0.1μg/ml LPS=48.17±1.75µm^2^, *P*<0.0001), comparable to swelling following 30 minutes OGD (MCC: 30 minutes OGD=36.5±0.36µm^2^ vs control *P*<0.0001, vs 0.1μg/ml LPS *P*>0.9999. MON: 30 minutes OGD=46.49±0.49µm^2^, vs control P<0.0001, vs 0.1μg/ml LPS *P*=0.8333). 30 minutes OGD + 0.1μg/ml LPS pre-application also caused soma swelling (MCC: 30 minutes OGD + 0.1μg/ml LPS=36.96±0.36µm^2^, *P*<0.0001. MON: 30 minutes OGD + 0.1μg/ml LPS=47.66±0.54µm^2^, *P*<0.0001), but not beyond 30 minutes OGD alone (MCC: *P*>0.9999. MON: *P*>0.9999).

Whilst low-grade inflammation did not elevate oligodendrocyte injury following ischemia, low-grade inflammation alone caused morphological signs of stress in oligodendrocytes that may have an impact on their ability to maintain healthy myelin during and following ischemia.

### Acute, low-dose LPS pre-application causes microglial activation, but no astrogliosis, without overt myelin and axonal damage

We next examined the response of other glial cells in white matter to examine their contribution to elevating white matter sensitivity to ischemia.

To examine the role of astrocytes, we used the astrocytic marker GFAP, which is acutely upregulated in the *ex vivo* MON in response to injury (Sup. Fig. 5.)^42^. There was no significant change in the number of GFAP+ cells (Control=152.7±5.87 cells/mm^2^, 0.1µg/ml LPS=145.8±5.13 cells/mm^2^, 30 minutes OGD=150.9±0.82 cells/mm^2^, 30 minutes OGD + 0.1µg/ml LPS=137.5±7.50 cells/mm^2^, *P*=0.2693) (Fig. 6Ai, B), fluorescence intensity (Control=44.88±3.55, 0.1μg/ml LPS=46.63±7.56, 30 minutes OGD=35.78±3.61, 30 minutes OGD + 0.1μg/ml LPS=37.50±4.37, *P*=0.3698) (Fig. 6Aii, B) or relative abundance of pixels positive for GFAP (Control=26.30±2.31%, 0.1μg/ml LPS=34.85±9.60%, 30 minutes OGD=35.78±2.61%, 30 minutes OGD + 0.1µg/ml LPS=37.50±4.37%, *P*=0.6461) (Fig. 6Aiii, B).

**Figure 6:**
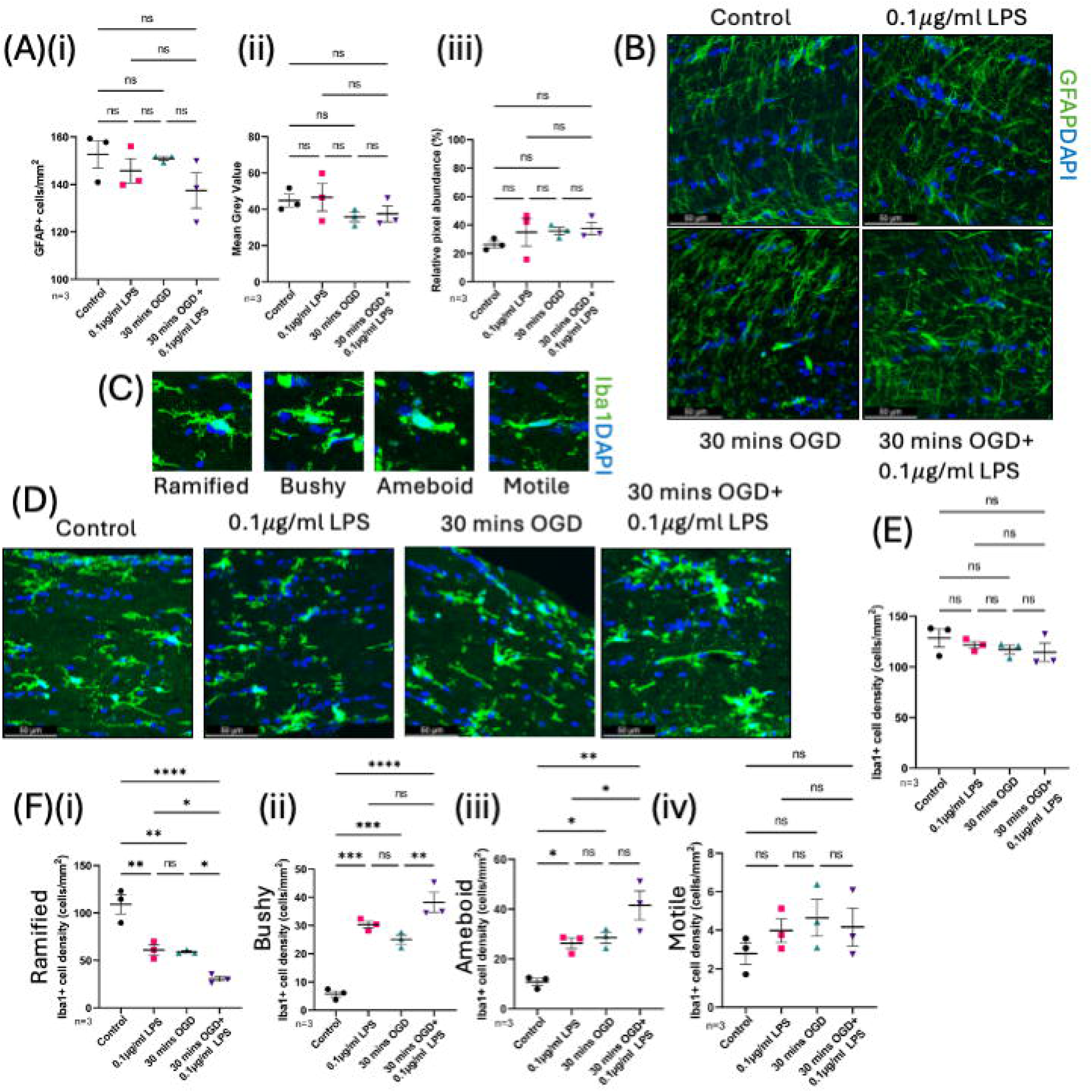
0.1μg/ml LPS causes no observable astrogliosis but subtle shifts in microglia morphology in the optic nerve. (A) **(i)** Density of GFAP+ astrocytes in the optic nerve following 0.1μg/ml LPS, 30 mins OGD or both (One-way ANOVA). **(ii)** Mean grey value of GFAP immunostaining in the optic nerve following 0.1μg/ml LPS, 30 mins OGD or both (One-way ANOVA). **(iii)** Relative abundance of GFAP+ pixels in the optic nerve following 0.1μg/ml LPS, 30 mins OGD or both (One-way ANOVA). **(B)** 40X Representative images of GFAP immunostaining in the optic nerve following 0.1μg/ml LPS, 30 mins OGD or both **(C)** Microglial morphologies in the optic nerve. **(D)** 40X Representative images of Iba1 immunostaining in the optic nerve following 0.1μg/ml LPS, 30 mins OGD or both. **(E)** Iba1+ cell density in the optic nerve following 40X following 0.1μg/ml LPS, 30 mins OGD or both (One-way ANOVA). **(F)** Density of ramified **(i)**, bushy **(ii)**, ameboid **(iii)** and motile **(iv)** microglia in the optic nerve following 0.1μg/ml LPS, 30 mins OGD or both (One-way ANOVA with Holm-Šídák’s post hoc multiple comparisons).

As the innate immune cells, and the major LPS-responsive glia in white matter, we looked to characterise the number and morphological response of microglia to 30 minutes OGD, 0.1µg/ml LPS or both, in the MON. Iba1+ microglial counts in the MON remained unchanged across all conditions (Control=128.4±8.86 cells/mm^2^, 0.1µg/ml LPS=121.7±3.41 cells/mm^2^, 30 minutes OGD=117.3±4.32 cells/mm^2^, 30 minutes OGD + 0.1µg/ml LPS=114.4±8.99 cells/mm^2^, *P*=0.5368) (Fig. 6D, E).

Next, we looked at the morphologies of microglia in the MON to identify whether they moved away from the ramified morphology indicative of resting, homeostatic microglia to morphologies associated with acute myelin injury^42^ (Fig. 6C, D, F). There were significant changes in all morphologies, apart from motile microglia (Ramified *P*<0.0001, Bushy *P*<0.0001, Ameboid *P*=0.0015, Motile *P*=0.4514) Microglia in control MONs were predominantly ramified (109.1±10.09 cells/mm^2^), with low numbers of bushy (5.76±1.09 cells/mm^2^), ameboid (10.82±1.43 cells/mm^2^) and motile (2.79±0.56 cells/mm^2^) microglia. Ramified microglia significantly decreased in MONs following 0.1μg/ml LPS (61.02±5.20 cells/mm^2^, *P=*0.0016), while bushy (30.34±1.21 cells/mm^2^, *P=*0.0008) and ameboid (26.32±2.13 cells/mm^2^ *P=*0.0283) morphologies increased. Further, there was no change in the number of motile microglia (3.98±0.61 cells/mm^2^). Following 30 minutes OGD, ramified microglia numbers were reduced (59.04±1.18 cells/mm^2^, *P*=0.0016), with increases in bushy (25.05±1.59 cells/mm^2^, *P*=0.0008) and ameboid microglia (28.54±2.13 cells/mm^2^, *P*=0.0283). Motile microglia numbers did not change (4.17±0.99 cells/mm^2^). Following combined 30 minutes OGD + 0.1μg/ml LPS pre-application, the number of ramified microglia further decreased (30.52±2.70 cells/mm^2^, vs. control *P*<0.0001, vs. 30 mins OGD *P*=0.0186). Bushy microglia numbers increased significantly (38.24±3.54 cells/mm^2^, *P*<0.0001 vs control, *P*=0.0186 vs 30 minutes OGD). Ameboid microglia numbers also rose (41.52±5.78 cells/mm^2^, *P*=0.0012 vs control), but not significantly from 30 minutes OGD alone (*P*=0.0052). The number of motile microglia remained unchanged (4.17±0.99 cells/mm^2^). This suggests that microglia are activated, even under low-grade inflammation, causing oligodendrocyte stress that may predispose them to elevated myelin loss.

### Microglial depletion removes the elevated sensitivity of white matter to OGD following acute, low-dose LPS pre-application

Given that microglial activation was observed, even under low-grade acute inflammation, we next looked to confirm whether microglial activation was responsible for elevating the sensitivity of white matter ischemia using pharmacological depletion with the CSF1R antagonist PLX5622.

21-day dietary administration of PLX5622 depleted microglia in MONs compared to mice fed a matched control diet (AIN-76A) and remaining microglia in depleted MONs did not exhibit morphological signs of activation (Sup. Fig. 6A, B).

Administration of AIN-76A control diet or PLX5622 combined with 0.1μg/ml LPS, 30 minutes OGD or both had significant effects on the size of the AUC of the CAP from the MON over the 110-hour recording period (*P*<0.0001) (Fig. 7A, Sup. Fig 7A). PLX5622 had no effect on the viability of MONs in control recordings or their sensitivity to 0.1μg/ml LPS across the recording period (Control, *P*>0.05 at all time points. 0.1μg/ml LPS, *P*>0.05 at all time points). Similarly, PLX5622 did not significantly alter recovery following 30 minutes OGD alone (*P*>0.05 at all timepoints). However, in AIN-76A MONs, combined 30 minutes OGD + 0.1μg/ml LPS pre-exposure significantly impaired recovery compared to 30 minutes OGD alone (*P*>0.05 from 62 to 110 mins, except at 79 mins; *P*=0.5133). This effect was abolished in PLX5622 MONs (*P*>0.05 at all timepoints).

**Figure 7:**
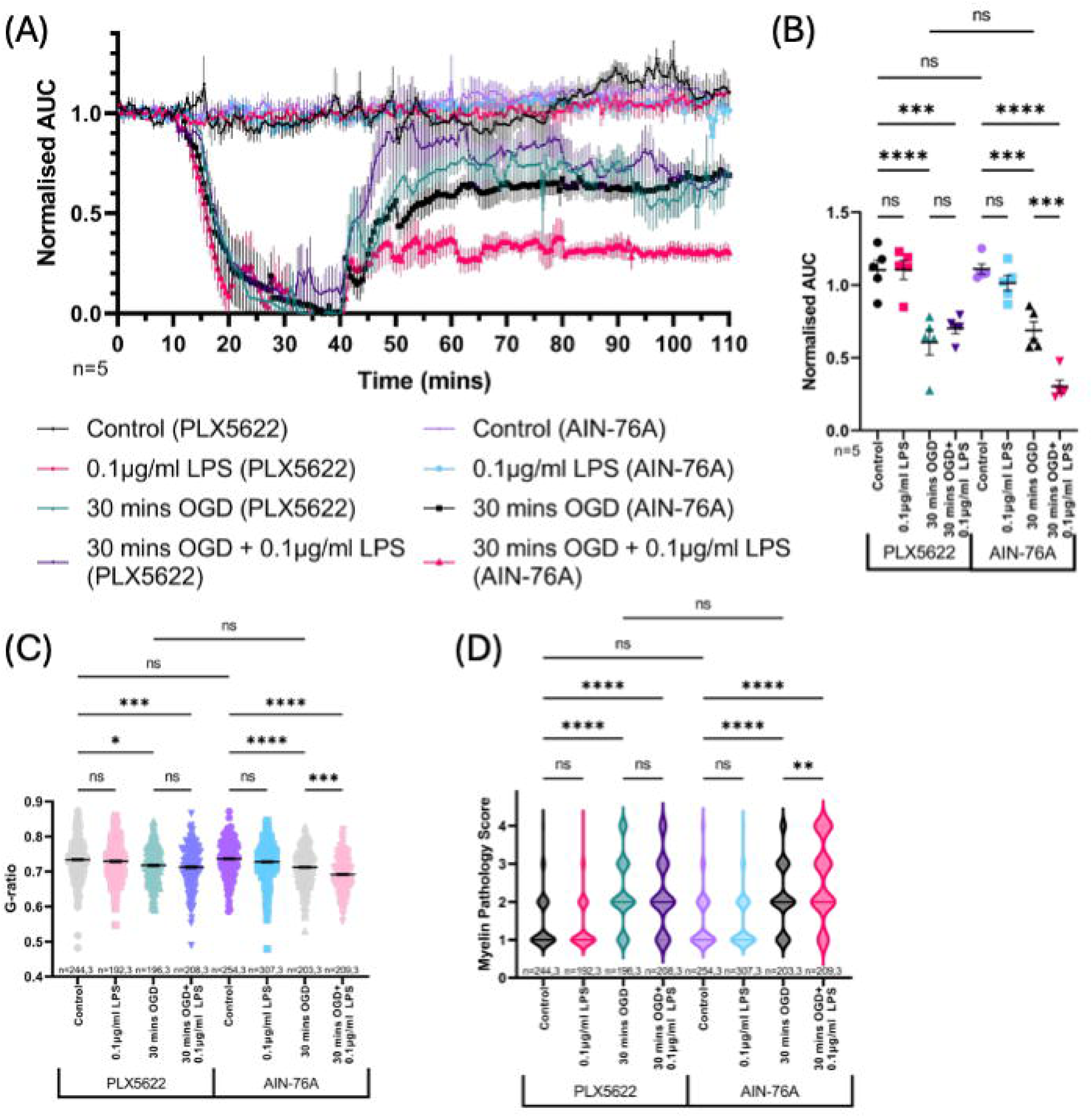
Microglial depletion reduces increased loss of optic nerve function and myelin injury following combined 30 minutes OGD and 0.1μg/ml LPS. **(A)** Normalised area under the curve of CAP recordings over time in control optic nerves and optic nerves treated with 0.1μg/ml LPS, 30 mins OGD or both, following 21-days PLX5622 diet or AIN-76A diet (Two-way repeated measures ANOVA with Holm-Šídák’s post hoc multiple comparisons). **(B)** Normalised area under curve of CAP recordings at 110 mins (endpoint) following 0.1μg/ml LPS, 30 mins OGD or both, following 21-days PLX5622 diet or AIN-76A diet (One-way ANOVA with Holm-Šídák’s post hoc multiple comparisons). **(C)** G-ratios of optic nerve axons following 0.1μg/ml LPS, 30 mins OGD or both following 21-days PLX5622 diet or AIN-76A diet (One-way ANOVA with Bonferroni’s post-hoc multiple comparison). **(D)** Myelin pathology scores of optic nerve myelin sheaths following 0.1μg/ml LPS, 30 mins OGD or both following 21-days PLX5622 diet or AIN-76A diet (Kruskal-Wallis test with Dunn’s post hoc multiple comparison).

Endpoint AUCs differed significantly across all groups (*P*<0.0001) (Fig. 7B). PLX5622 had no effect on the AUC in control MONs (PLX6522=1.10±0.07, AIN-67A=1.11±0.04, *P*=0.9998), following 0.1μg/ml LPS (PLX5622=1.11±0.07, AIN-76A=1.01±0.05, *P=*0.9271) or following 30 minutes OGD (PLX5622=0.61±0.09, AIN-76A=0.69±0.06, *P*=0.9271). In AIN-76A MONs, 30 minutes OGD + 0.1μg/ml LPS pre-exposure significantly reduced AUC compared to AIN-76A alone (30 minutes OGD=0.69±0.06, 30 minutes OGD + 0.1μg/ml LPS=0.30±0.04, *P*=0.0007). This reduction was absent in PLX5622 MONs (30 minutes OGD=0.61±0.09, 30 minutes OGD + 0.1μg/ml LPS=0.70±0.04, *P*=0.9271).

We assessed G-ratios and myelin pathology in MONs from mice fed AIN-76A or PLX5622 and exposed to 0.1μg/ml LPS, 30 minutes OGD or both (Fig. 7, Sup. Fig. 7B). G-ratios differed significantly across groups (P<0.0001) (Fig. 7C). PLX5622 had no significant effect on the G-ratio in control MONs (AIN-76A=0.74±0.003, PLX5622=0.73±0.004, *P*>0.9999), following 0.1μg/ml LPS (AIN-76A=0.73±0.003, PLX5622=0.73±0.004, *P*>0.9999). or following 30 minutes OGD (AIN-76A=0.71±0.003, PLX5622=0.72±0.003, *P*>0.9999). In AIN-76A MONs, combined 30 minutes OGD + 0.1μg/ml LPS pre-application significantly reduced the G-ratio compared to 30 minutes OGD alone (30 minutes OGD=0.71±0.003, 30 minutes OGD + 0.1μg/ml LPS=0.69±0.004, *P*=0.0006), while in PLX5622 MONs, this reduction in G-ratio was lost (30 minutes OGD=0.72±0.004, 30 minutes OGD + 0.1μg/ml LPS=0.71±0.004, *P*>0.9999).

Myelin pathology scores differed significantly across conditions (*P*<0.0001) (Fig. 7D). PLX5622 had no effect on myelin pathology scores in the control MON (AIN-76A=1.52±0.05, PLX5622=1.44±0.44, *P*>0.9999), following 0.1μg/ml LPS (AIN-76A=1.46±0.04, PLX5622=1.37±0.04, *P*>0.9999) or 30 minutes OGD (AIN-76A=2.15±0.05, PLX5622=2.27±0.07, *P*<0.9999). In MONs from AIN-76A-fed mice, the myelin pathology score significantly increased between MONs following 30 minutes OGD or 30 minutes OGD + 0.1μg/ml LPS pre-application (30 minutes OGD=2.15±0.05, 30 minutes OGD + 0.1μg/ml LPS=2.60±0.07, *P*<0.0013). Whereas, in MONs from mice fed PLX5622, there was no significant difference in the myelin pathology score following 30 minutes OGD or 30 minutes OGD + 0.1μg/ml LPS pre-application (30 minutes OGD=2.27±0.07, 30 minutes OGD + 0.1μg/ml LPS=2.16±0.06, *P*<0.9999).

Therefore, we show that depleting microglia in the MON removes the heightened sensitivity to ischemia caused by low-grade inflammation.

### Minocycline removes the elevated functional loss of white matter following OGD and acute, low-dose LPS pre-application

Following identification that the depletion of microglia eliminates the reduction of white matter sensitivity to ischemia, we next examined whether manipulation of microglia with the anti-inflammatory drug minocycline could also eliminate this elevated sensitivity.

0.1μg/ml LPS, 30 minutes OGD and combined treatment, both with and without 10μM minocycline, had significant effects on the AUC of the CAP over time (*P*<0.0001) (Fig. 8A B). 10μM minocycline did not affect the AUC in control MONs (*P*>0.05 at all timepoints) or MONs treated with 0.1μg/ml LPS (*P*>0.05 at all timepoints). Recovery following 30 minutes OGD was unchanged with or without 10μM minocycline (*P*>0.05 at all timepoints). In untreated MONs, combined 30 minutes OGD + 0.1μg/ml LPS pre-application significantly impaired recovery compared to 30 minutes alone (*P*<0.05 from 63.5 to 110 mins). This difference was lost in MONs treated with 10μM minocycline (*P*>0.05 at all timepoints).

**Figure 8:**
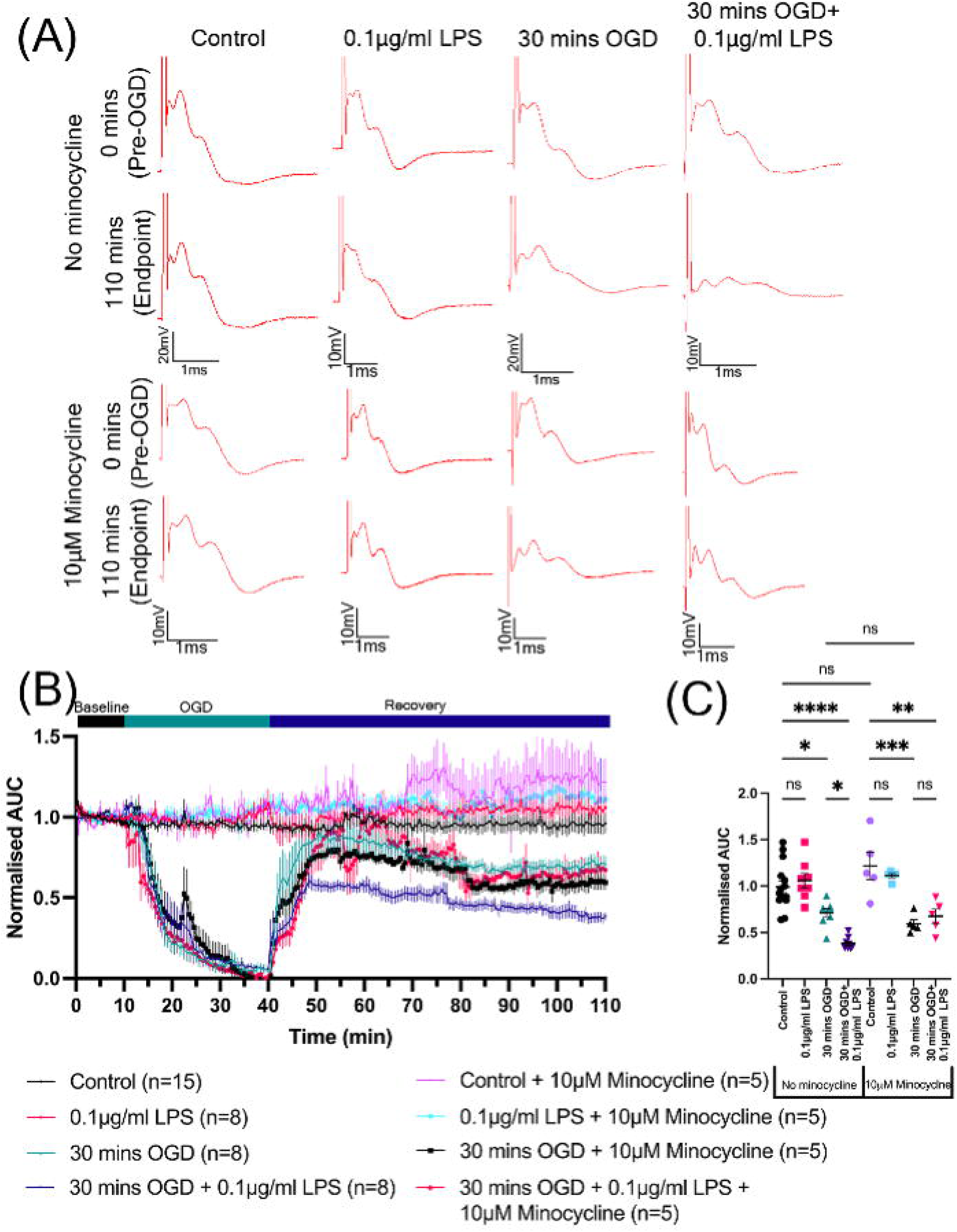
Minocycline reduces the increased loss optic nerve function following combined 30 minutes OGD and 0.1μg/ml LPS. **(A)** Representative CAP traces at 0 mins (baseline, pre-30 mins OGD) and 110 mins (endpoint, post-70 mins recovery). **(B)** Normalised area under the curve of CAP recordings over time in control optic nerves and optic nerves treated with 0.1μg/ml LPS, 30 mins OGD or both alongside 10μM minocycline (Two-way repeated measures ANOVA with Holm-Šídák’s post hoc multiple comparisons). **(C)** Normalised area under curve of CAP recordings at 110 mins (endpoint) following 0.1μg/ml LPS, 30 mins OGD or both alongside 10μM minocycline (One-way ANOVA with Holm-Šídák’s post hoc multiple comparisons).

Endpoint AUCs differed across conditions (*P*<0.0001) (Fig. 8C). There was no effect of minocycline in control MONs (No minocycline=0.99±0.06, 10μM minocycline=1.22±0.15, *P*=0.2517), response to 0.1μg/ml LPS (No minocycline=1.06±0.08, 10μM minocycline=1.11±0.02, *P*=0.9217) or 30 minutes OGD (No minocycline=0.71±0.05, 10uM minocycline=0.60±0.04, *P*=0.8852). Without minocycline, there was a significant difference in the endpoint AUC following 30 minutes OGD and 30 minutes OGD + 0.1μg/ml LPS pre-application (30 minutes OGD=0.71±0.05, 30 minutes OGD + 0.1μg/ml LPS=0.38±0.02, *P*=0.0211). This significant difference was lost following treatment with 10μM minocycline (30 minutes OGD=0.60±0.04, 30 minutes OGD+0.1μg/ml LPS=0.68±0.08, *P*=0.9271).

This therefore shows that targeting white matter with drugs that reduce microglial activation can remove white matter sensitivity to ischemia, indicating minocycline prevents inflammation-induced exacerbation of acute ischemic injury.

## Discussion

Given the link between acute inflammatory diseases and ischemic brain injury^5–14^, we investigated whether low-grade white matter inflammation affects susceptibility to ischemic damage using isolated white matter tracts. Functional and cellular analyses revealed that while acute, low-grade inflammation does not directly damage white matter, it increases injury under ischemic conditions.

### Acute, low-grade inflammation elevates the ischemic sensitivity of white matter

White matter employs intrinsic stress-tolerance mechanisms, including glutamate clearance by oligodendrocytes and astrocytes preventing excitotoxicity^44–46^ and ROS detoxification by oligodendrocytic and myelinic peroxisomes, histidine dipeptides and antioxidant enzymes^47–50^, maintaining function under stress. Acute, low-dose exposure to the TLR-4 agonist LPS caused no axonal or myelinic damage or functional changes, indicating tolerance of acute, low-grade inflammatory conditions; established mechanisms of injury appear sub-critical, allowing for continued function during low-grade inflammatory challenges. Despite this, oligodendrocytes show signs of stress such as somatic and mitochondrial swelling following low-grade inflammation ^43^. Oligodendrocytes induce stress responses to protect themselves against injury, such as the integrated stress response^51–53^, but this may reduce resilience to subsequent injury. Following ischemia, there was reduced functional recovery of white matter that coincided with an elevated level of axonal and myelinic damage when preparations were pre-inflamed. This suggests coincidental inflammation expends oligodendrocytes stress tolerance, impairing their ability to maintain healthy myelin during ischemia. Oligodendrocytes, and by extension their myelin sheaths, have a limited capacity to handle stressors. Acute, low-grade inflammation likely uses a certain amount of stress tolerance capacity, therefore exceeding the threshold for damage quicker and resulting in increased damage, when acute, low-grade inflammation occurs alongside ischemia, elevating myelin damage.

Elevated myelin damage was reflected in elevated damage to axons. While axons remained largely intact after acute, low-grade inflammation, co-exposure led to more severe ischemic damage. This may result from elevated damage directly, for example oxidative damage^54^, or as secondary effects of glial dysfunction. Axons depend heavily on glia- especially myelin- for metabolic support and injury protection^55–57^. Loss of oligodendrocytic support is a key driver of axonal degeneration in demyelinating diseases^58^. Consequently, impaired glial function and worsened myelin integrity may exacerbate mitochondrial stress and neurofilament breakdown in axons, compounding the effects of acute, low-grade inflammation and ischemia in white matter.

### Microglia: central but targetable mediators of elevated ischemic sensitivity

Microglia actively sense and respond to CNS injury and are key players in both ischemic and inflammatory conditions. Their activation is a hallmark of these diseases^59,60^, potentially positioning them at the interface of the interaction of ischemic and low-grade inflammatory injury.

Following acute, low-grade inflammation, microglia shifted to a subtle pro-inflammatory state, adopting morphologies associated with acute myelin injury^42^. This was unsurprising given their central role in responding to inflammatory stimuli^61,62^. This morphological shift coincided with increased oligodendrocyte stress measures. Activated microglia release pro-inflammatory cytokines such as interferons^51,52,63,64^, glutamate and ROS^23,65–69^, all of which can trigger oligodendrocyte stress. Thus, this pro-inflammatory shift likely impairs oligodendrocytes’ ability to maintain healthy myelin during ischemia.

Microglia shift to a pro-inflammatory state during acute, low-grade inflammation, likely driving increased white matter sensitivity to ischemia, consistent with their role in other conditioning paradigms^70^. To confirm this, we depleted microglia using the CSF1R antagonist PLX5622^71,72^. Microglial-depleted white matter was electrophysiologically viable and showed no differences in functional response or myelin injury following LPS treatment or ischemia. However, microglial depletion eliminated the increased ischemic damage following combined acute, low-grade inflammation and ischemia in measures of function and myelin injury, showing microglia are central to the elevated sensitivity to white matter ischemia. This likely reflects reduced oligodendrocyte stress due to fewer microglial-derived cytokines, ROS and extracellular glutamate, all known contributors to oligodendrocyte injury^23,65–69^.

As microglia appeared central to increased ischemic sensitivity during acute, low-grade inflammation, we asked whether they could be therapeutically targeted. Minocycline, a clinically available drug known to dampen microglial responses^73–82^, including reducing ROS and cytokine release and promoting anti-inflammatory phenotypes^83–85^, was evaluated for its ability to prevent microglial-mediated elevation of ischemic injury. Similar to microglial depletion, minocycline eliminated the elevated functional sensitivity to ischemia, confirming that pharmacologically reducing microglial activation mitigates the increase in ischemic sensitivity. Although minocycline shows mixed translational results^86–89^, we provide proof-of-concept that targeting microglia may prevent excessive white matter injury during ischemia combined with inflammation, with potential implications for at-risk patients.

### White matter sensitization: a mechanism behind elevated risk

We show that acute, low-grade inflammation in white matter can elevate white matter sensitivity to ischemia, providing mechanistic insight into why patients with a higher inflammatory burden are more vulnerable to ischemic brain injuries. Acute, low-grade inflammation may create a sub-critical neuroinflammatory state, predisposing white matter to damage during otherwise asymptomatic or transient ischemic events^90–93^, such as silent infarcts or transient ischemic attacks. Combined with acute, low-grade inflammation, these minor events could trigger or amplify injury, leading to irreversible ischemic damage that would not occur without neuroinflammation.

We modelled this epidemiological link in a simplified *ex vivo* setting using a low-grade LPS model to mimic aspects of *in vivo* inflammation. By applying a pro-inflammatory mediator directly to central white matter lacking the blood-brain barrier (BBB), we mimic conditions where CNS immune privilege is compromised. Peripheral inflammation elevates cytokines and chemokines in the CNS through BBB crossing or signalling through meningeal and endothelial cells^94^. LPS itself can cross the BBB via lipoprotein receptors^95^, and infections often increase BBB permeability^96,97^. In white matter, TLR-4 activation by LPS causes upregulation of pro-inflammatory molecules^98^, replicating the clinical inflammatory milieu. Therefore, low LPS concentrations may replicate the low-grade inflammatory state seen during human infection.

This work may inform public health by clarifying infection-related risk factors for ischemic brain injury. With no effective treatments for ischemic brain injury, prevention through managing the risks of inflammatory conditions, like that shown by the management of other risk factors such as hypertension^99^, remains key. Understanding how infections elevate stroke risk may also strengthen public messaging; for instance, linking oral pathogens to stroke risk reinforces the importance of good oral health.

## Conclusions

Here we show that white matter tolerates acute, low-grade inflammation via intrinsic protective mechanisms, however, responds to subsequent ischemic insults with elevated axonal and myelin damage, highlighting the limited stress capacity of oligodendrocytes. Microglia drive the increased sensitivity of white matter to ischemia following acute, low-grade inflammation. Both microglial depletion and minocycline prevent this effect, highlighting microglia as key therapeutic targets. Given the sensitization effects we observe here, it suggests why patients with a higher inflammatory burden face greater risk of ischemic brain injury, where sub-critical inflammatory states may amplify damage during minor or transient ischemic events, highlighting inflammation as both a risk factor and therapeutic target.

## Acknowledgements

Thanks to Professor David Parkinson, University of Plymouth for providing the PLX5622 diet used in these experiments and Mr. Waldemar Woznica and colleagues at the University of Plymouth Biological Services Unit for the care of animals used in this study.

## Supplementary material

Supplementary material is available at *Brain* online.

---

**Supplementary figure 1:**
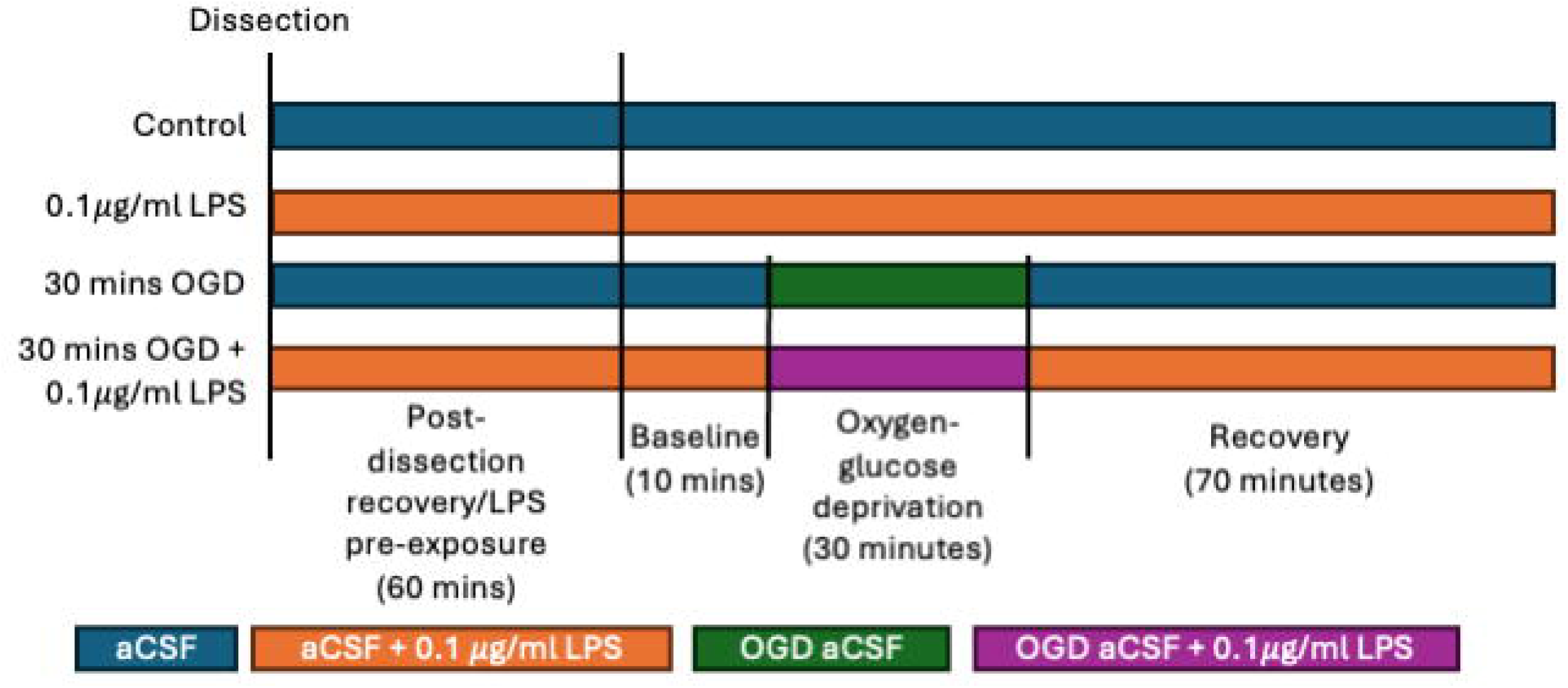
Experimental time course of *ex vivo* tissue preparations.

**Supplementary figure 2:**
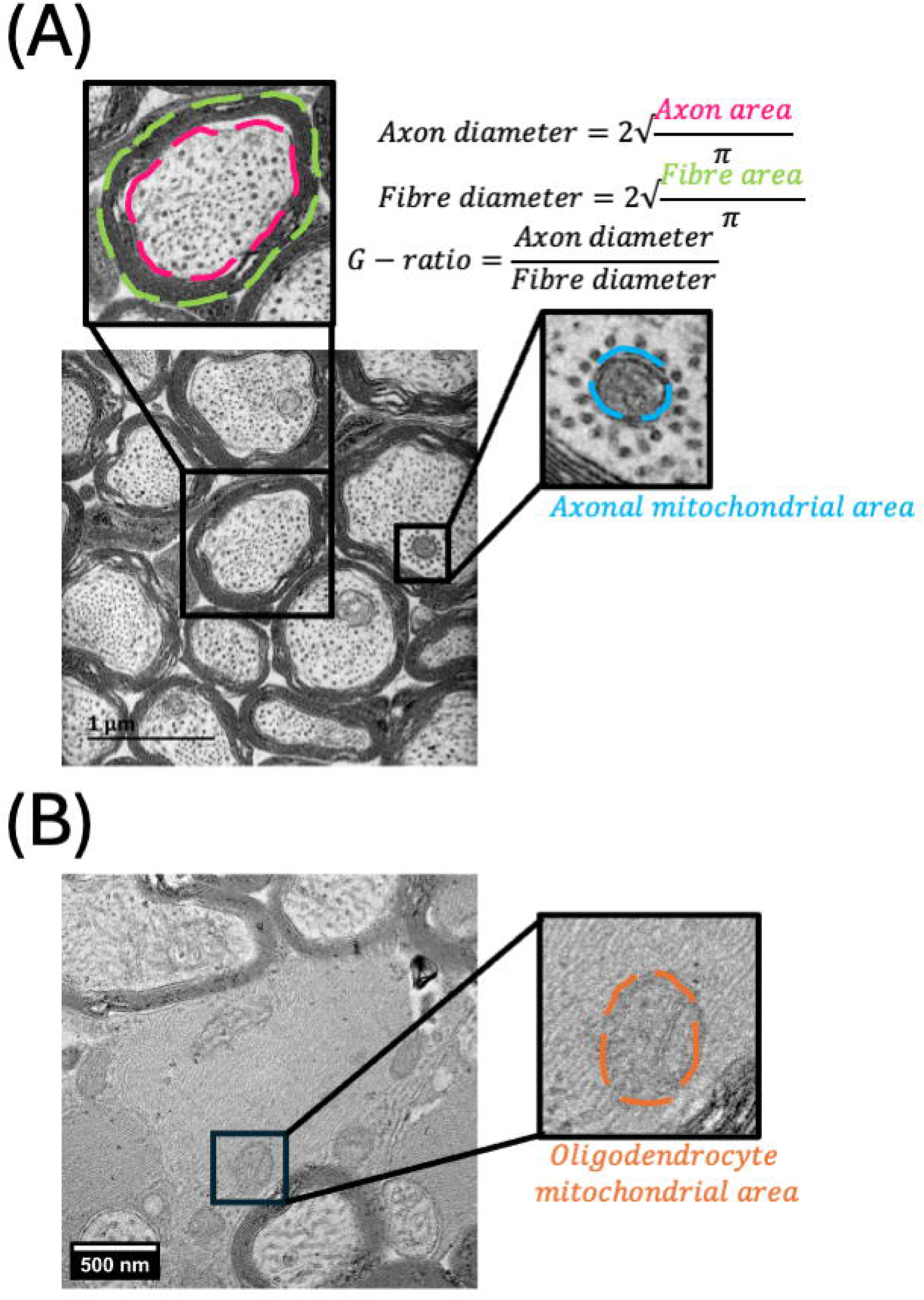
Electron microscopy quantification of optic nerve myelinated axons and oligodendrocytes. **(A)** Fibre and axonal diameter and axonal mitochondrial area measurements and G-ratio calculation. **(B)** Oligodendrocyte mitochondrial area measurement.

**Supplementary figure 3:**
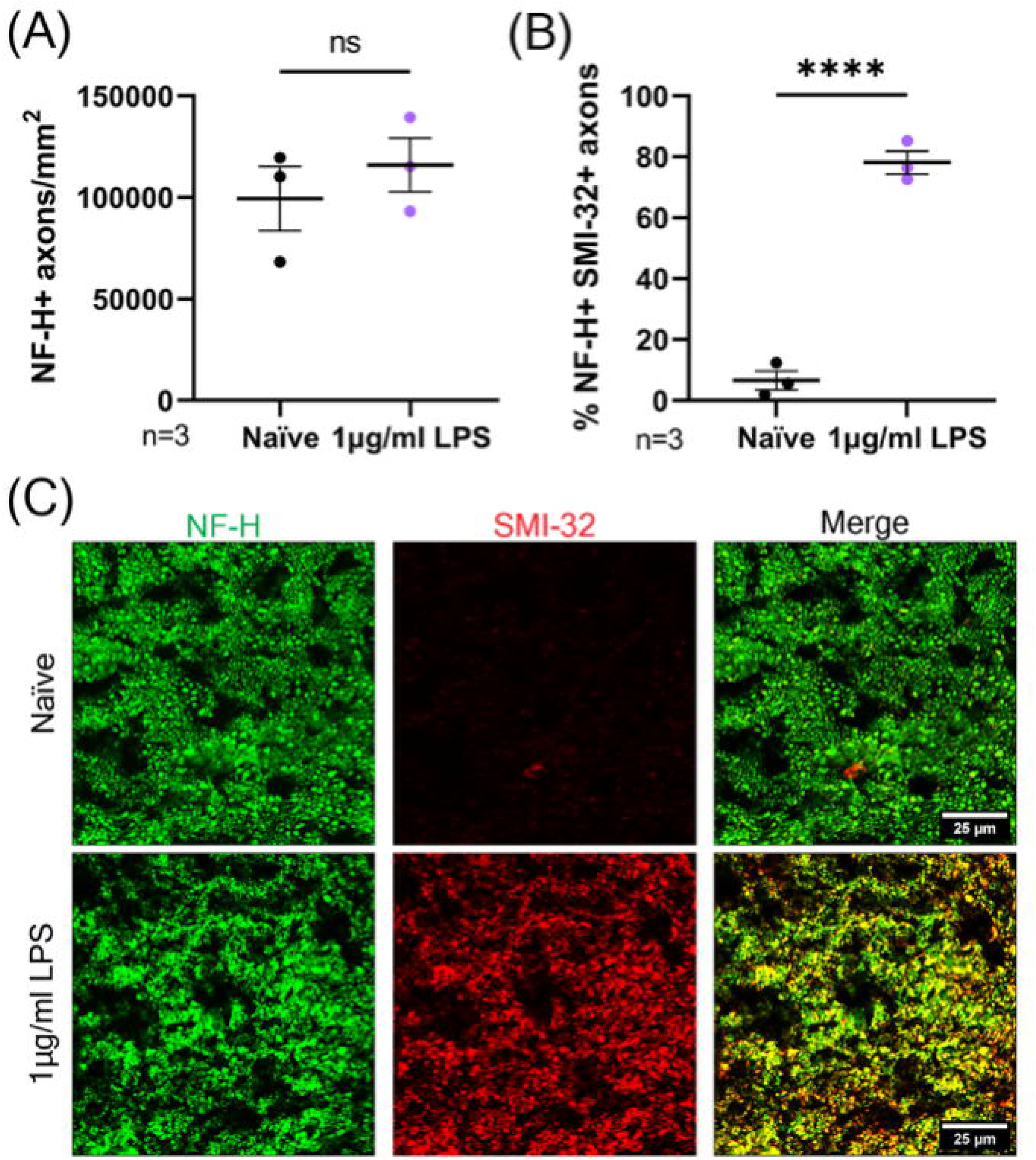
1μg/ml LPS causes an increase in dephosphorylated neurofilament in the optic nerve. **(A)** The density of NF-H+ axons in the optic nerve following 120 mins treatment with 1μg/ml LPS (unpaired t-test). **(B)** The percentage of NF-H+ axons positive for SMI-32 following 120 mins treatment with 1μg/ml LPS (unpaired t-test). **(C)** 63X representative images of optic nerve axons immunostained for NH-H and SMI-32.

**Supplementary figure 4:**
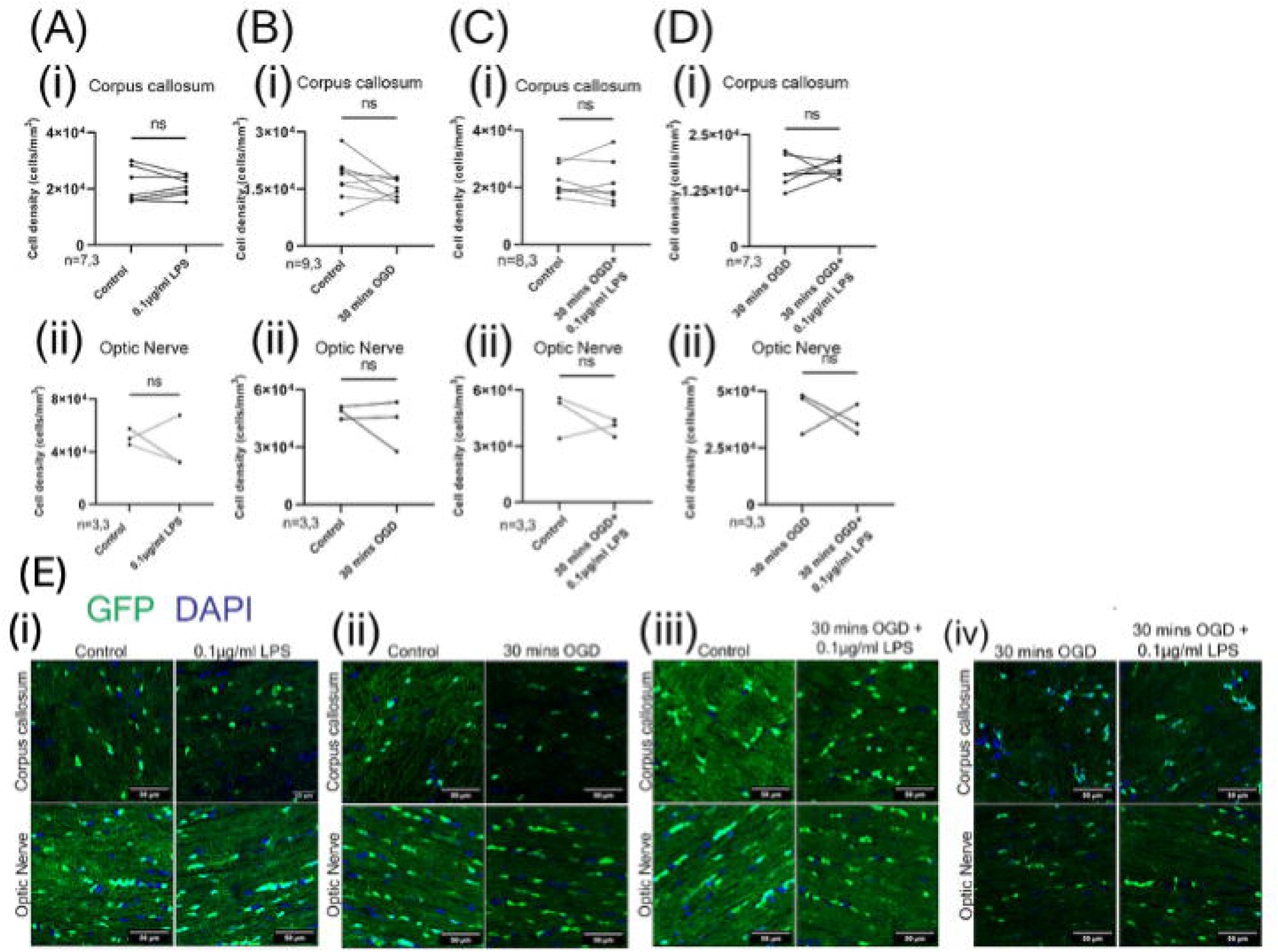
0.1μg/ml LPS, 30 minutes OGD or both causes no change in the density of oligodendrocytes in the corpus callosum or optic nerve (A-D) The density of GFP+ oligodendrocytes in the corpus callosum **(i)** and the optic nerve **(ii)** following 0.1μg/ml LPS, 30 mins OGD or both (paired t-test). **(E)** 40X representative images of the corpus callosum and optic nerve following 0.1μg/ml LPS, 30 mins OGD or combined.

**Supplementary figure 5:**
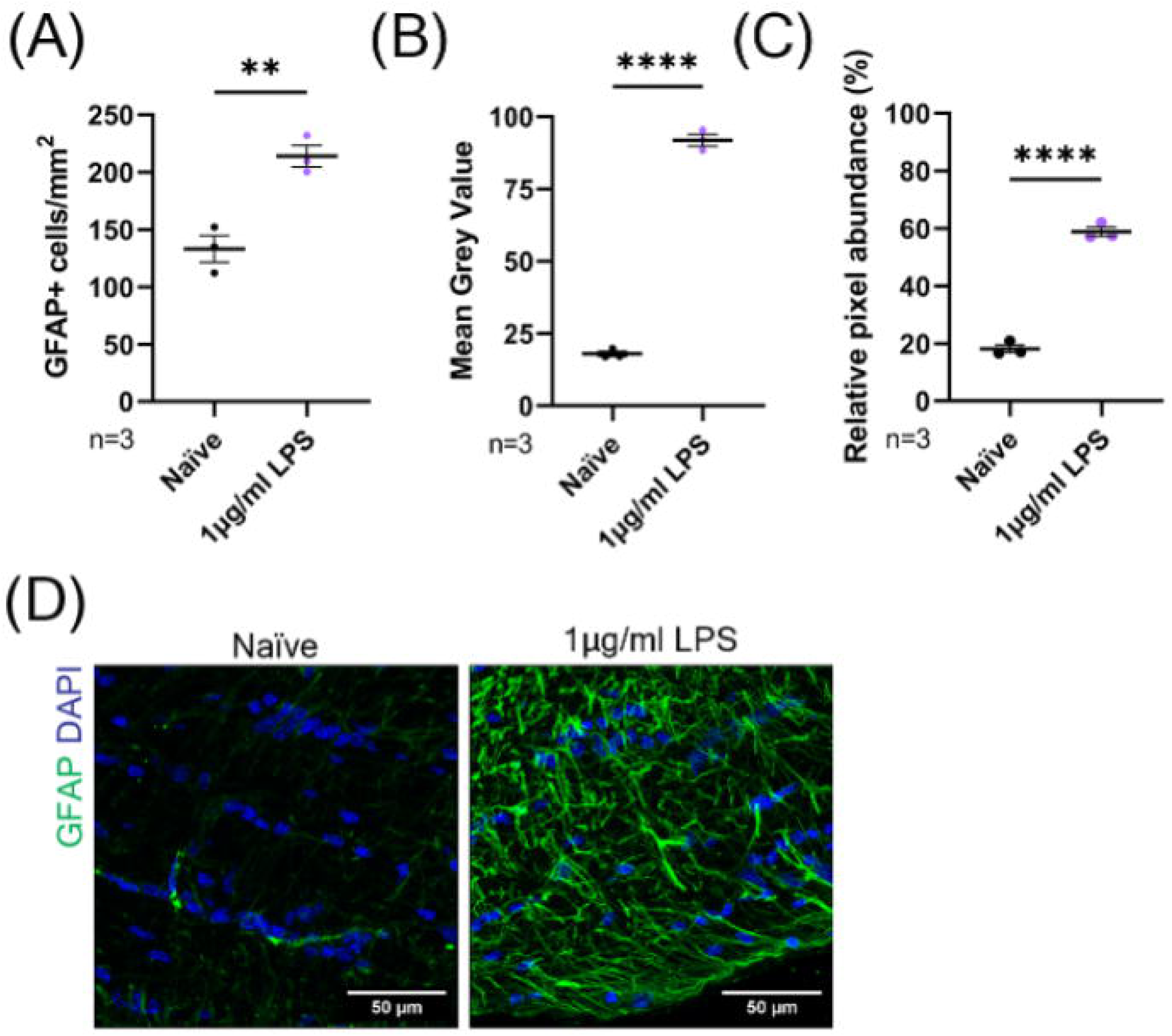
1μg/ml LPS causes astrocytosis in the optic nerve. **(A)** The number of GFAP+ cells in the optic nerve following 120 mins treatment with 1μg/ml LPS (Unpaired t-test). **(B)** The mean grey value of GFAP immunostaining in the optic nerve following 120 min treatment with 1μg/ml LPS (Unpaired t-test). **(C)** Relative pixel abundance of GFAP in the optic nerve following 120 min treatment with 1μg/ml LPS (Unpaired t-test). **(D)** 40X representative images of optic nerves following control and 1μg/ml LPS treatment for 120 mins.

**Supplementary figure 6:**
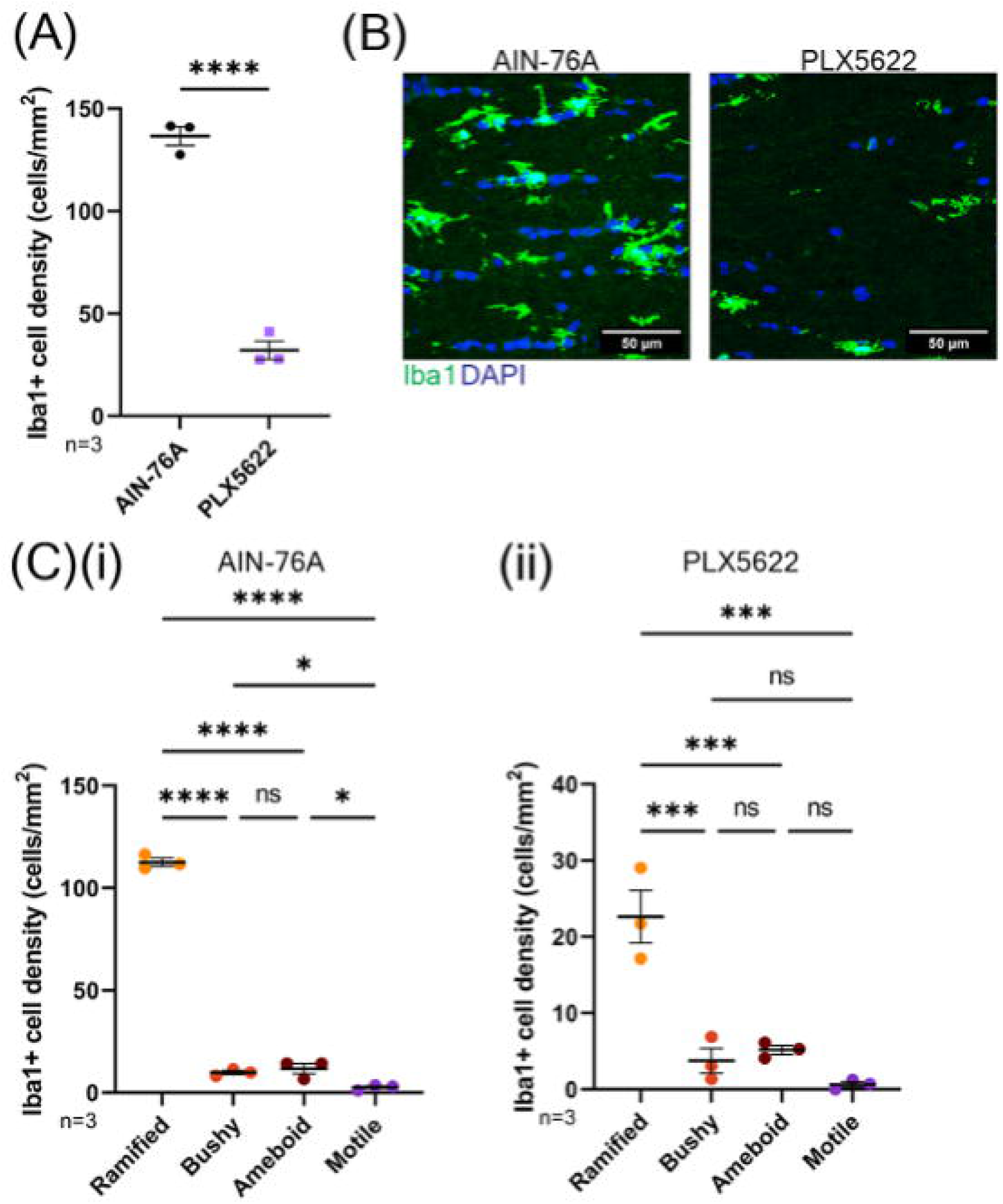
21 days treatment with PLX5622 depletes microglia in the optic nerve. **(A)** Density of Iba1+ cells in the optic nerve following 21 days PLX5622 treatment (Unpaired t-test). **(B)** 40X representative images of the optic nerve following 21 days PLX5622 treatment. **(C)** Morphologies of Iba1+ microglia in the optic nerve following AIN-76A diet **(i)** and PLX5622 treatment **(ii)** for 21 days combined (One-way ANOVA with Holm-Šídák’s post hoc multiple comparisons).

**Supplementary figure 7:**
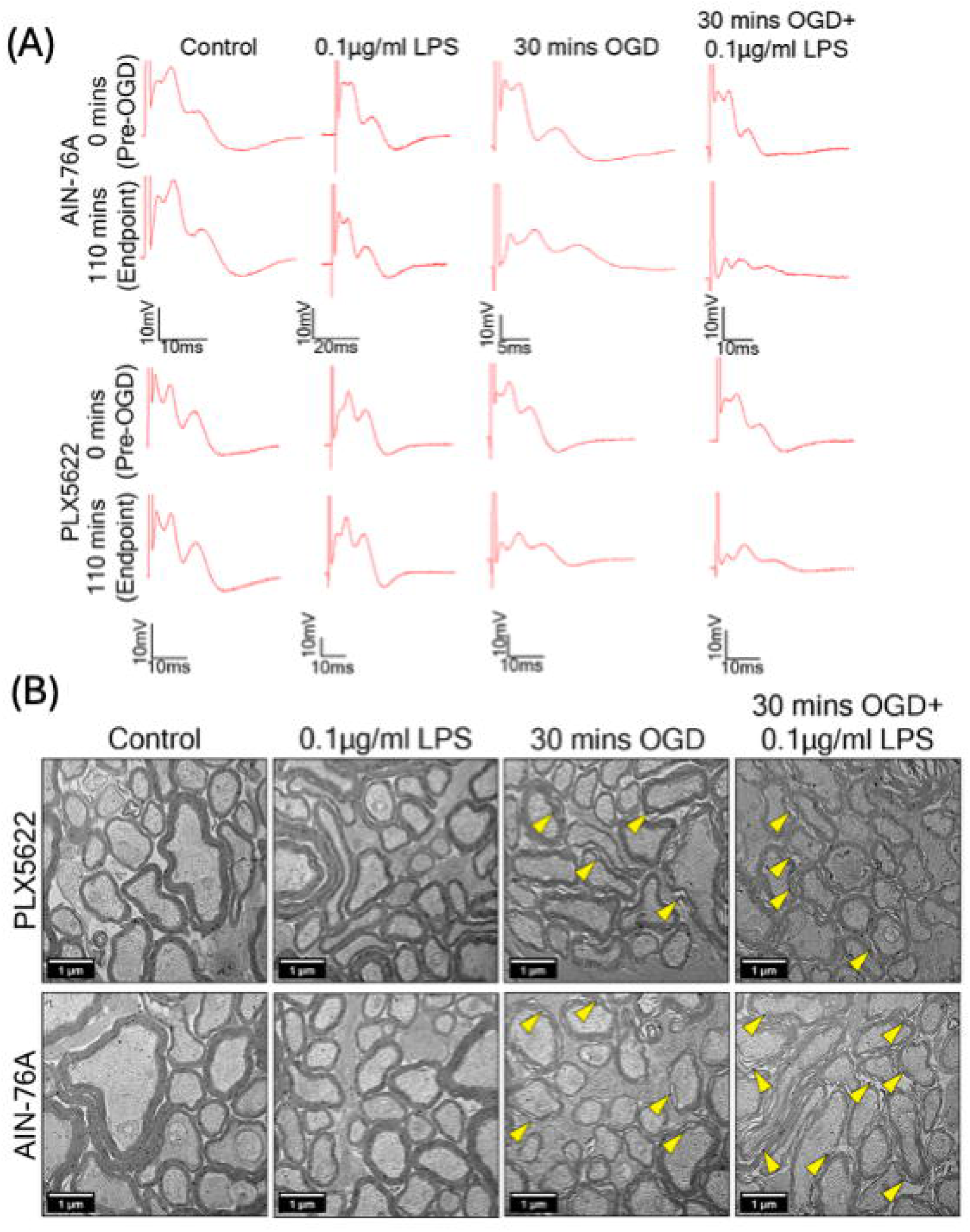
Electrophysiological and ultrastructural observations following microglial depletion with PLX5622. **(A)** Representative CAP traces at 0 mins (baseline, pre-30 mins OGD), 40 mins (post-30 mins OGD), and 110 mins (endpoint, post-70 mins recovery). **(B)** 10,000X representative micrographs of the optic nerve; focal decompaction sites indicated by yellow arrows.

